# The immune synapses reveal aberrant functions of CD8 T cells during chronic HIV infection

**DOI:** 10.1101/2022.01.13.476200

**Authors:** Nadia Anikeeva, Maria Steblyanko, Leticia Kuri-Cervantes, Marcus Buggert, Michael R. Betts, Yuri Sykulev

**Author notes:** Department of Medicine Huddinge, Karolinska Institutet, Karolinska University Hospital Huddinge (Marcus Buggert).

## Abstract

It is well-established that chronic HIV infection causes persistent low-grade inflammation that induces premature aging of the immune system in HIV patient including senescence of memory and effector CD8 T cells. To uncover the reasons of gradually diminished potency of CD8 T cells from chronically HIV infected people, we have analyzed cellular morphology and dynamics of the synaptic interface followed exposure of peripheral polyclonal CD8 T cells at various differentiation stages to planar lipid bilayers. The above parameters were linked to pattern of degranulation that determines efficiency of CD8 T cells cytolytic response. We found a large fraction of naive T cells from HIV infected people developing mature synapses and demonstrating focused degranulation, a signature of a differentiated T cells. Further differentiation of aberrant naïve T cells leads to development of anomalous effector T cells undermining their capacity to control HIV and other viruses that could be contained otherwise.

## INTRODUCTION

Essential role of cytotoxic CD8 T cells in anti-viral immunity is well established ^1–4^. Particularly, depleting CD8+ T cells in primates resulted in complete failure to suppress initial burst of SIV replication suggesting that CD8 T cells make critical contribution to control of viremia shortly after infection and during chronic phase of the infection ^5, 6^. Recognition of infected cells by the T cells result in triggering of T-cell receptor (TCR)- and integrin-mediated signaling that initiates the formation of contact area which transforms into highly organized structure called immunological synapse (IS) ^7, 8^. The formation of this structure contributes to the coordinated delivery and release of cytolytic granules to target cells, which trigger apoptosis in targeted cells ^9, 10^. IS at the T cell/target cell interface represents complex 3D structure that is very challenging to study using microscopy because the area is not flat, highly dynamic, and include multiple components, the role of each of them is difficult to ascertain. Glass-supported planar lipid bilayers that mimic the surface of target cells provides unmatched opportunity to analyze T cell contact area, especially dynamics of the molecular events at the interface ^10, 11^. It has been well established that two proteins, ICAM-1 and pMHC ligands, displayed on lipid bilayer is sufficient to induce the formation of immune synapse by T cells exposed to bilayers ^12, 13^, very similar to that observed at T cell/target cell interface ^7, 14^. This permits us to evaluate differences in the structure and dynamic of synaptic interface formed by individual polyclonal T cells with various phenotypes and functional activities.

Available data suggest that HIV infection could lead to a global defect in T cells including CD8 T cells ^15, 16^. Our previous studies of human T cell clones that are derived from peripheral blood of infected people, particularly those with HIV infection, have demonstrated that efficiency of cytolytic activity exercised by cytotoxic T lymphocytes (CTL) is linked to kinetics of TCR-mediated Ca^2+^ signaling and the structure and stability of IS formed by CTL ^10, 17^. However, whether the parameters of synaptic interface of primary T cells varies at different stages of differentiation has not be studied. This further solidifies the need to investigating the structure and dynamics of IS of polyclonal T cells from HIV infected and uninfected people using planar lipid bilayers ^18, 19^.

With this in mind, we have examined synaptic interface formation by peripheral CD8 T cells at various differentiation stages from HIV-infected and uninfected people. The comparison revealed significant differences in the dynamic, structure, and stability of the interface and also exhibited notable dissimilarity in the pattern and kinetics of the cell’s degranulation. Unexpectedly, we have found that significant fraction of naïve CD8 T cells from HIV-infected people form mature synapses and a large fraction of these cells degranulate. Our findings suggest that chronic inflammation during HIV infection mediates changes in the ability of T-cells to form synaptic interface and, consequently, their functional activity.

## RESULTS

### The structure of T-cell/bilayer interfaces produced by PBMC-derived CD8 T cells from uninfected and HIV-infected people

We exposed polyclonal CD8 T cells derived from uninfected and HIV-infected people to planar lipid bilayers that display anti-CD3 antibodies and ICAM-1. Majority of the T cells adhere to the bilayers and form T cell/bilayer interfaces. Size of adhesion area, dynamic, structure and stability of the interfaces were examined. Regardless of donor status, analysis of these parameters revealed four categories of CD8 T cells interacting with the bilayers. Only a fraction of cells established “mature synapse” (**Fig. 1A, Movies 1A and B**) that resembled a classical bull-eye structure and adhesion area similar to those observed previously for cloned CD8 T cells ^10, 13^. Another fraction of round cells that wobbled over the bilayer were not able to spread and have a small adhesion area (**Fig. 1B, Movies 2A and B)**. These cells formed “focal synapses” that constitute of accumulated TCR/CD3 without peripheral ring junction containing adhesion molecules but revealed poor and transient ICAM-1 accumulation ^20^. Still others formed a very unstable dynamic synapses or “kinapses” ^21^ showing migratory morphology with asymmetric adhesion area (**Fig. 1C**, **Movies 3A and B**). Yet another category of T cells had a large adhesion area and formed lamellipodium. These cells often did not achieve complete segregation of CD3/TCR and LFA-1, and the T cell/bilayer interfaces appeared as multifocal synapses (**Fig. 1D, Movies 4A and B**).

**Figure 1.**
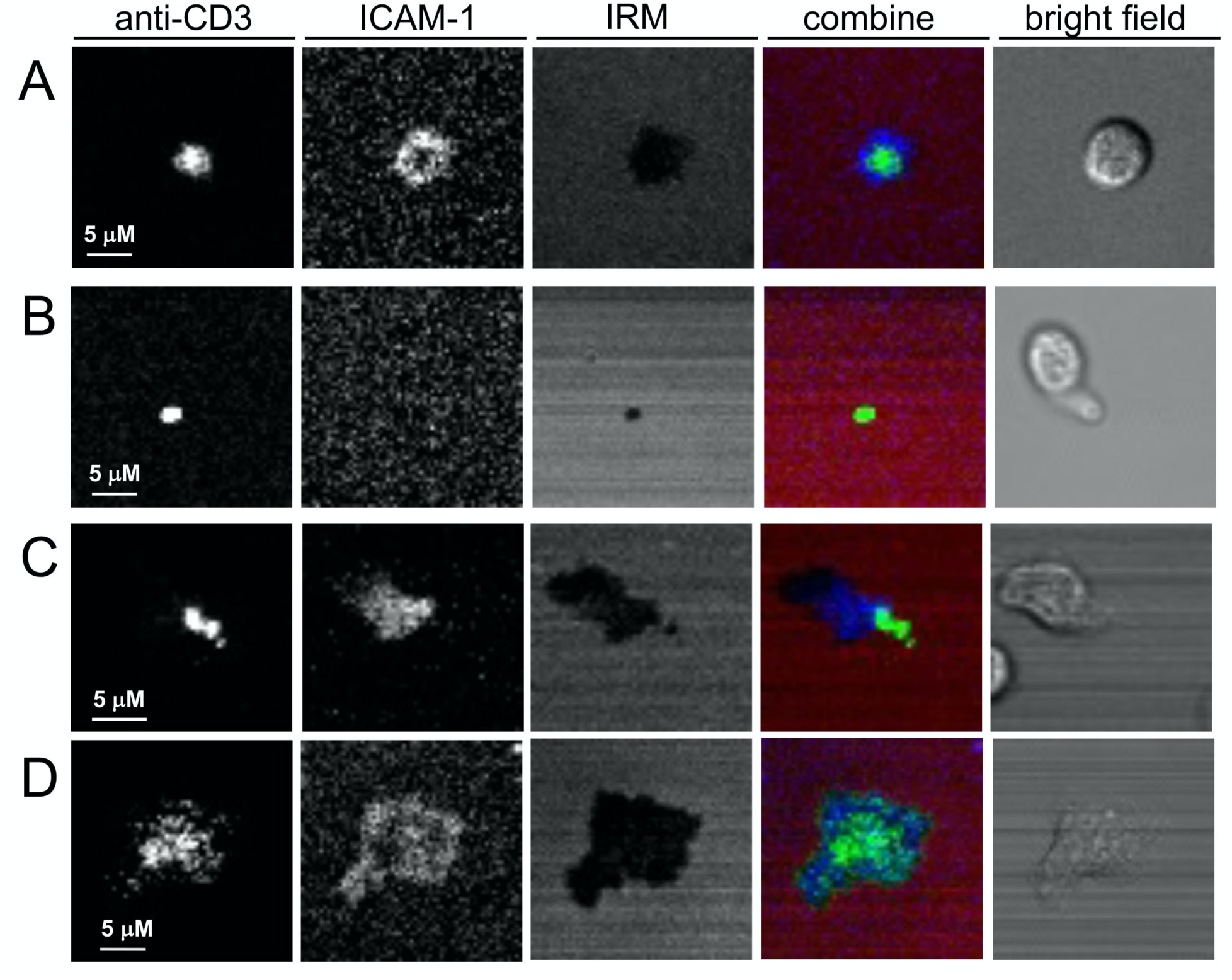
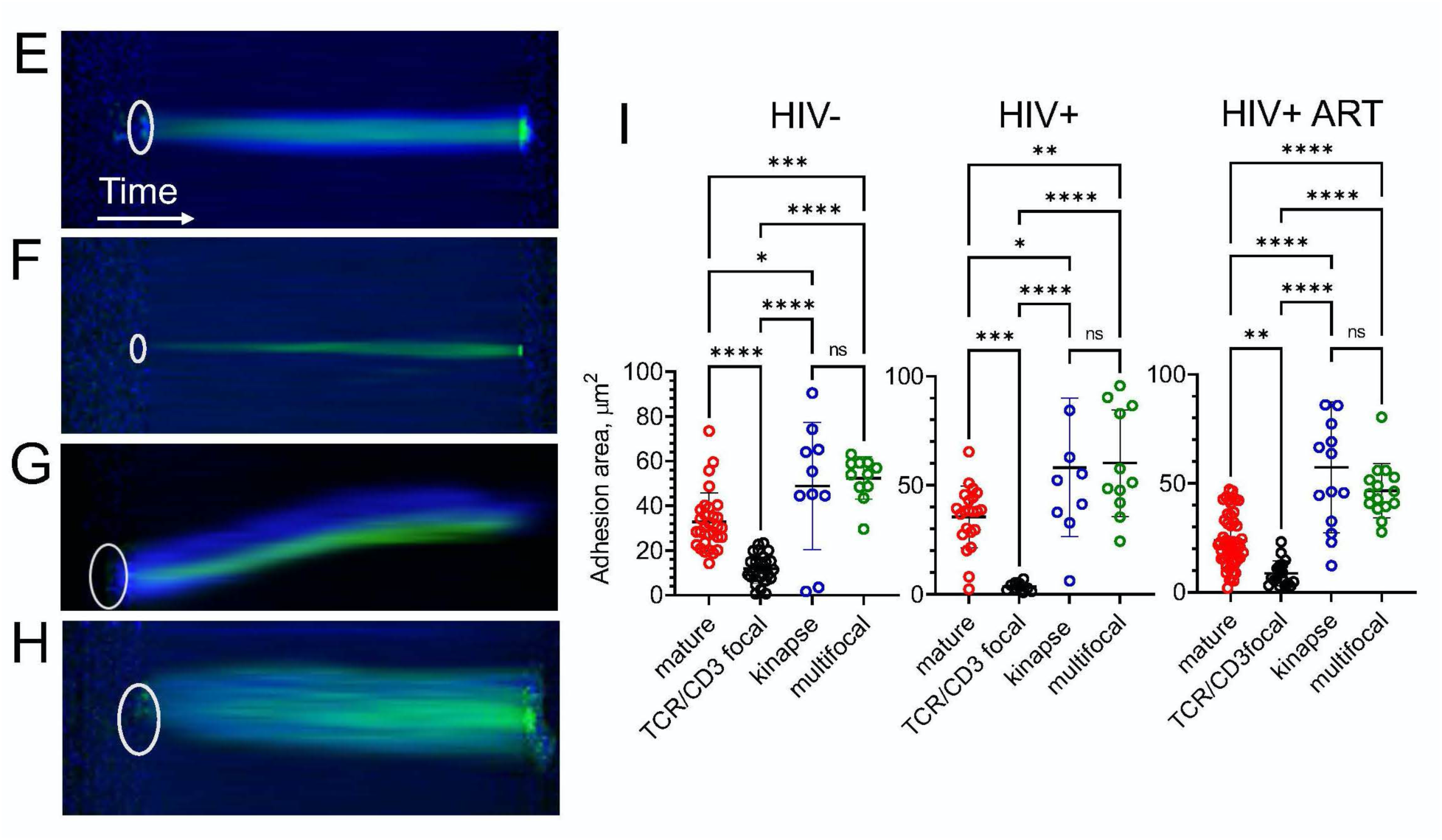
Human polyclonal CD8 T cells establish diverse synaptic interfaces with lipid bilayers that display labeled anti-CD3 antibody and ICAM-1 ligands. CD8 T cells were isolated from PBMC using negative magnetic sorting. Representative images of T cell-bilayer interface are shown: mature synapse (**A**), TCR/CD3 focal interface (**B)**, kinapse (**C)** and multifocal synapse (**D**). The images were taken by confocal microscope at 30 min after initial cell-bilayer contact. Representative kymograms show temporal changes in spatial position of ICAM-1 and anti-CD3 antibodies: mature synapse (**E**), TCR/CD3 focal interface (**F)**, kinapse (**G)** and multifocal synapse (**H**). The images were taken by confocal microscope every 2 min during 30 min. White oval indicates the position of the cell/bilayer interface at the initial contact of T cell with a bilayer surface, and time axis indicates temporal changes in the spatial position of ICAM-1 and CD3 locations. ICAM-1 is blue, anti-CD3 is green. IRM images show contacts of CD8 T cells with the bilayer surface as dark area on light background (red on overlay images). Scale bars represent 5 μm. Real-time images are available in Movies 1-4. Variations in adhesion area of different synaptic interfaces formed by CD8 T cells from uninfected, HIV-infected and treated with ART people are depicted in panel **I.** Adhesion area of the cells was calculated from IRM images (n≥ 10). Data are shown as mean ± SD; *p<0.05, **p<0.01, ***p<0.001, ****p<0.0001 as established by one-way ANOVA using Tukey post hoc test. At least two independent experiments for each condition were analyzed.

Changes of the interface structure in time for each category of the T cells are clearly evident from kimographs demonstrating temporal variations in spatial position of accumulated TCR/CD3 and LFA-1/ICAM-1 moieties (**Fig. 1E, F, G, H**).

We have also compared size of adhesion area and extent of LFA-1/ICAM-1 accumulation for the T cells derived from either HIV-infected or uninfected people as well as from patients treated with ART (**Fig. 1I**). We have found statistically significant difference in the average size of adhesion area between all T cell categories except T cells that form kinapses and lamellipodia. This was regardless of donors’ status. However, extent of ICAM-1 accumulation was similar for each T cell category, but the T cells that form focal synapses (**Supplemental Figure 1**).

The observed differences in the dynamics of cytolytic synapse formation, the synapses stability, the size and shape of adhesion area, and ICAM-1 accumulation are most likely linked to distinct stages of T cell differentiation and also reflect quality of T-cell functioning. With this in mind, we went on to purify and compare various subsets of CD8 T cells from PBMC of uninfected and HIV-infected individuals.

### The ability to accumulate ICAM by CD8+ T cells depends on their differentiation stage and donor status

Using magnetic sorting, we purified CD27+, CD27-CD45RO+ and CD27-CD45RO-CD8 T cells from PBMC of uninfected and infected donors **(Supplemental Figure 2)** and analyzed the ability of T cells to accumulate ICAM-1 at the T cell/bilayer interface followed ICAM-1 engagement by LFA-1 on the T cell surface (**Fig. 2**). For HIV uninfected donors, fraction of T cells capable to accumulate of ICAM-1 was dependent on the stage of T cell differentiation and progressively increased from 40% in CD27+ T cells to 75% observed in CD27-CD45RO-T cells (**Fig. 2A**). In marked contrast, more than 85% of CD27+ T cells derived from HIV-positive people were capable to accumulate ICAM-1 (**Fig. 2B**). There was no difference in the ability to accumulate ICAM-1 by CD27-CD45RO+ and CD27-CD45RO-T cells derived from HIV-infected people. ART treatment did not change the percentage of T cells showing ICAM-1 accumulation. (**Fig. 2C**).

**Figure 2.**
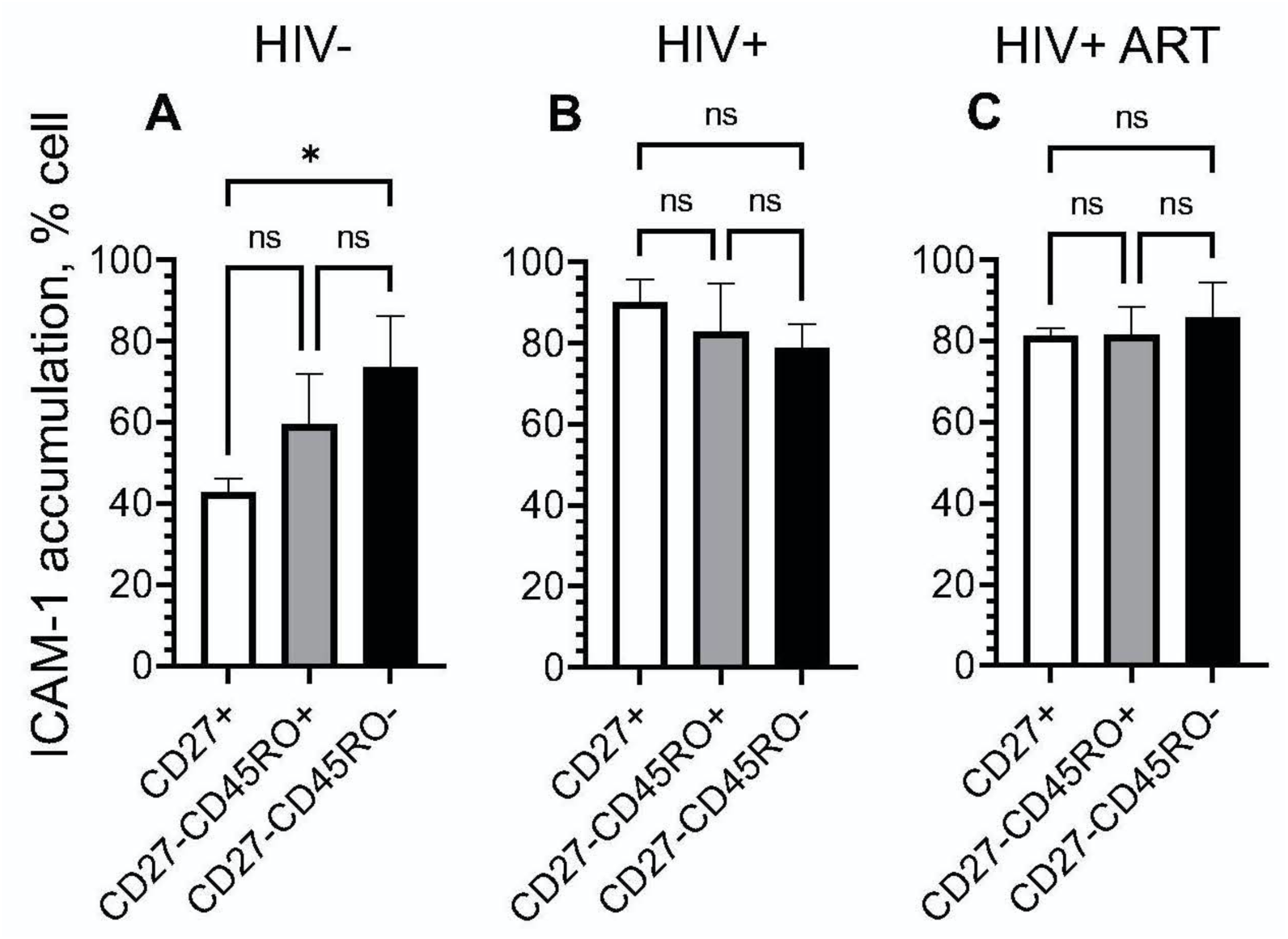
Fraction of CD8 T cells accumulating ICAM-1 at synaptic interface depends on T cell differentiation stage and donors’ status. CD8 T cells derived from PBMC of HIV- (**A**) or chronic HIV+ (**B**) or ART-treated HIV+ (**C**) individuals were subdivided on CD27+, CD27- CD45RO+ and CD27-CD45RO- subsets by magnetic sorting. Isolated T cell subsets were exposed to bilayers containing labeled anti-CD3 and ICAM-1 molecules, and ICAM-1 accumulation were observed at the interface for 30 minutes. The data represent mean values (±SD) of three independent experiment for each donor group with different infection status. Statistical significance within a subject group was determine using the one-way ANOVA using Tukey post hoc test; *p<0.05.

### Comparison of LFA-1 expression on CD8 T cells at various stages of differentiation

Accumulation and segregation of fluorescent labeled ICAM-1 incorporated into the lipid bilayers is mediated by its engagement with LFA-1 molecules on T cells. Polyclonal population of CD8 T cells regardless of the donors’ status contains cells that present either high or low level of LFA-1 expression (**Fig. 3A**). LFA-1^high^ population expressed about 3-5-times higher number of LFA-1 molecules per cell compared with LFA-1^low^ populations (**Supplemental Figure 3**). Analysis of LFA-1^high^ population for the T cells at various differentiation stages derived from HIV-infected or uninfected donors is shown in **Fig. 3B**. In HIV-uninfected individuals, only a small fraction of LFA-1^high^ was detected in CD27+ T cells. In marked contrast, significantly higher fraction LFA-1^high^ (≍80%) was observed in CD27+ T cells from HIV-infected people. High level of LFA-1 expression on majority of CD27+ T cells derived from HIV-infected people resembles antigen-experienced T cells. The presence of LFA-1^high^ T cells as well as number of LFA-1 molecules per cell in populations of CD27-CD45RO+ and CD27-CD45RO- cells were very similar and were not depended on donors’ status (**Fig. 3B** and **Supplemental Figure 4**). ART-treatment did not result in significant change of the fraction of CD27+ cells presenting high level of LFA-1 (**Fig. 3B**).

**Figure 3.**
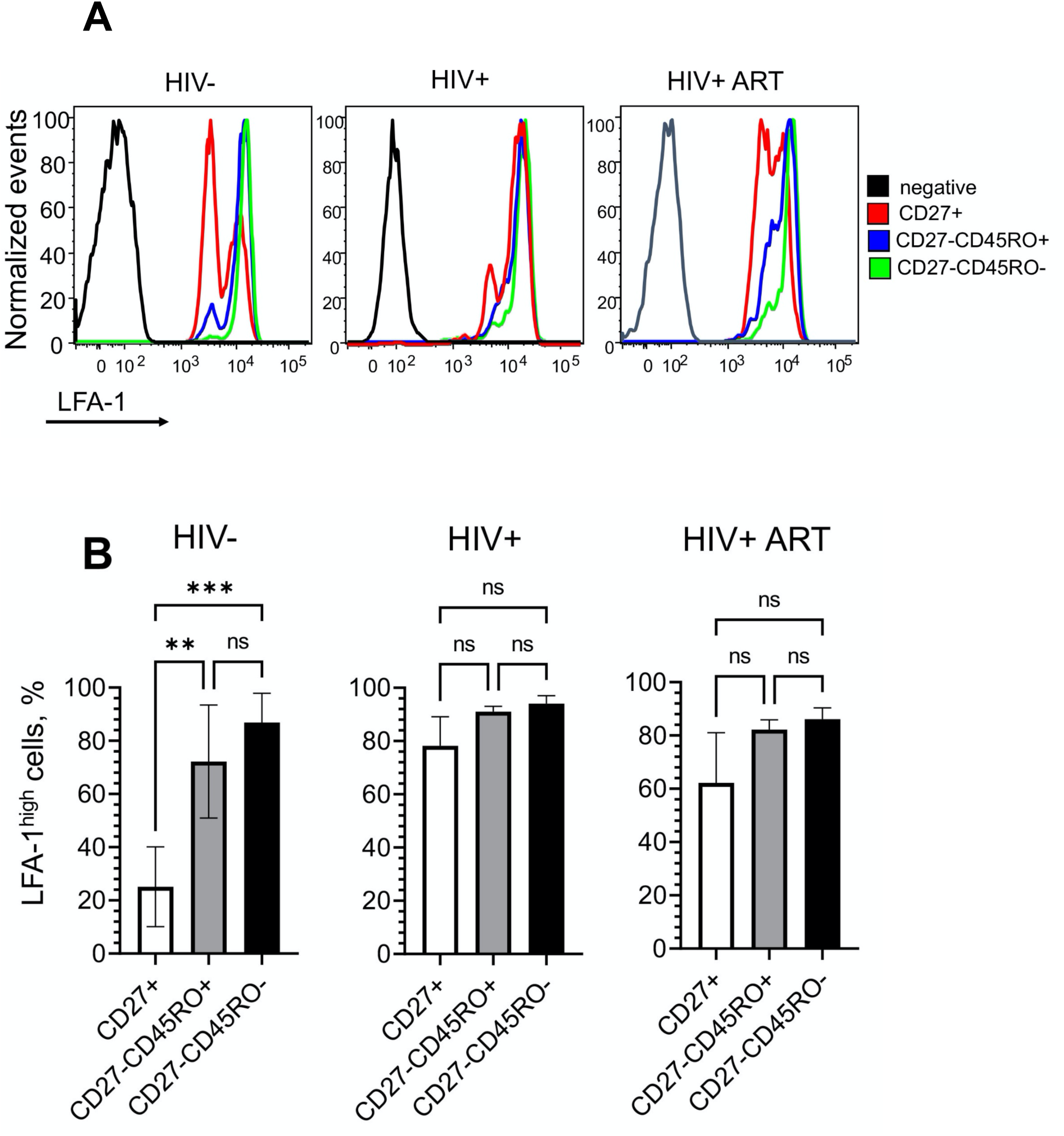
CD27+CD8 T cells from donors with chronic HIV infection contains significantly higher fraction of cells with upregulated LFA-1 expression, and ART treatment only partially return the size of the fraction to a normal level. The CD8 T cells subsets were isolated by magnetic sorting, stained with anti-LFA antibodies and analyzed by Flow Cytometry. (**A**) Representative flow histograms showing LFA-1 expression by CD27+, CD27-CD45RO+ and CD27-CD45RO-CD8 T cells derived from HIV- (left), chronic HIV+ (middle) and ART-treated HIV+ (right) donors. (**B**) Frequency of CD8 T cell with high LFA-1 expression within each subset for HIV- (left), chronic HIV+ (middle) and ART-treated HIV+ (right) individuals. For each condition, bar represent standard deviation of the mean of three independent experiments; **p<0.01, ***p<0.001 were determined by one-way ANOVA with Tukey post hoc test.

These data are consistent with the difference in the extent of ICAM-1 accumulation at in the interface of CD27+ CD8 T cells from uninfected and infected people (**Fig. 2**). This suggests that the LFA-1 engagement by ICAM-1 contributes significantly to the differences of the T cells to accumulate ICAM-1 at T cell/bilayers interfaces and to establish mature synapses.

### Early differentiated CD8 T cells from HIV+ donors revealed enhanced ability to form mature synapses

We determined proportions of T cells at various differentiation stages that possessed distinct structures of T cell/bilayer interfaces. A large fraction of CD27+ T cells derived from infected patients form mature synapses as opposed to those from uninfected donors (**Fig. 4A, B** and **Supplemental Figure 5**). Only a small fraction of the T cells with this phenotype was observed in HIV uninfected donors, while majority of the T cells concentrated TCR/CD3 without ICAM-1 accumulation establishing focal synapses (**Fig. 4A, B** and **Supplemental Figure 6**). The T cells from HIV+ individuals treated with ART exhibited a large fraction of T cells that capable to establish kinases, while the ability of these T cells to form mature synapse was similar to those derived from uninfected donors (**Fig. 4A, B** and **Supplemental Figure 5**). At the same time, T cells at more advanced differentiation stages (CD27-CD45RO+ and CD27-CD45RO-) did not show significant differences in the ability to establish synaptic interfaces of different kinds. The exception was CD27-CD45RO-T cells from HIV+ donors either treated or untreated with ART that revealed enhanced ability to form multifocal synapses and lamellipodia (**Fig. 4A** and **Supplemental Figure 7**).

**Figure 4.**
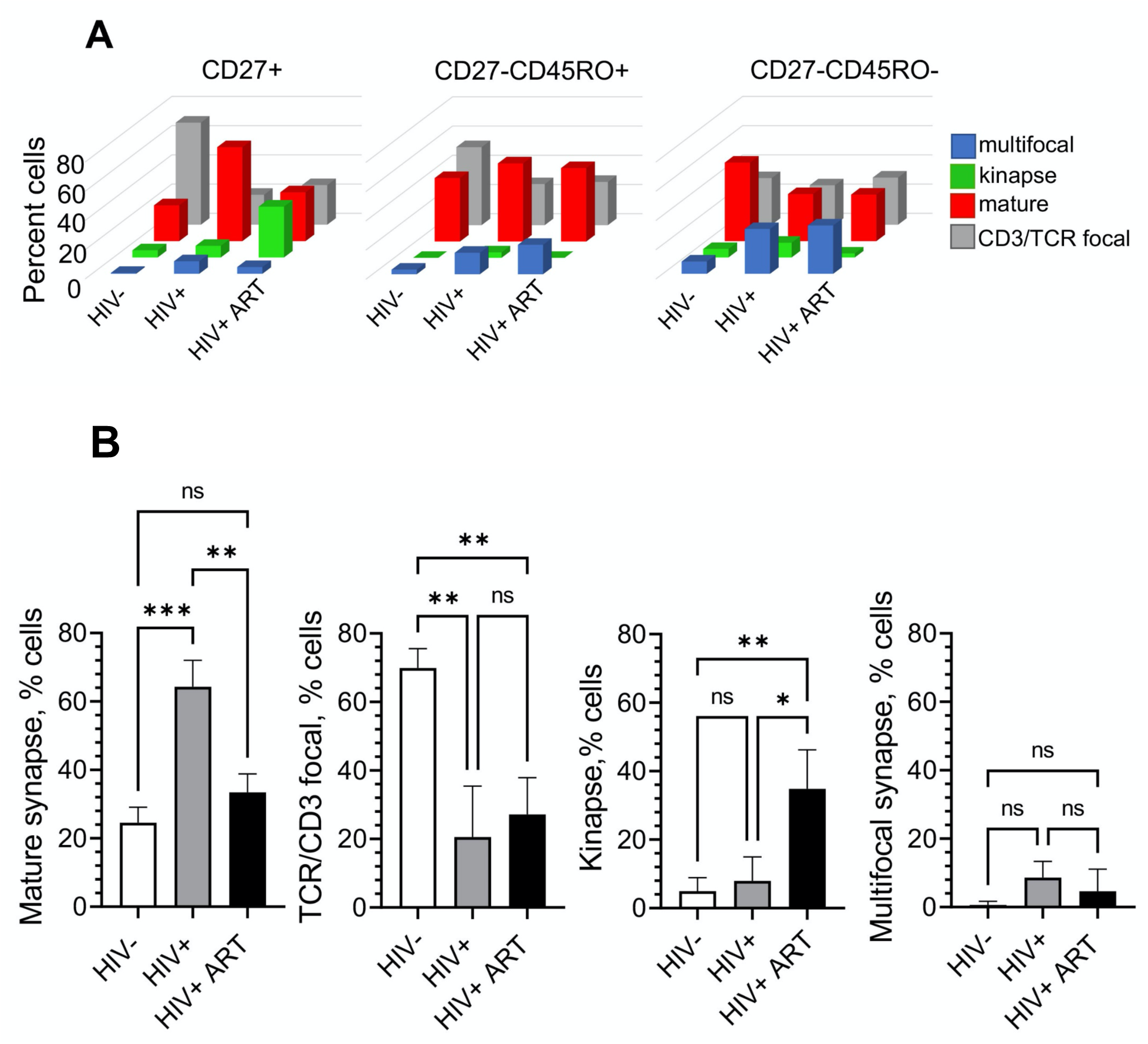
Dissimilarity of the synaptic interfaces established by polyclonal human CD8 T cells from HIV-, chronic HIV+ and ART-treated HIV+ individuals are most pronounce at early stage of differentiation. CD27+, CD27-CD45RO+ and CD27-CD45RO-subsets of CD8 T cells from HIV-, chronic HIV+ and ART-treated HIV+ individuals were isolated by magnetic sorting and loaded on bilayers containing anti-CD3 antibodies and ICAM-1. The formation of synaptic interfaces was observed for 30 minutes by confocal microscopy. (**A**) 3D plot represent frequency of CD27+ (left), CD27-CD45RO+ (middle) and CD27-CD45RO- (right) CD8 T subsets that form mature synapses, TCR/D3 focal interfaces, multifocal synapses and kinapses at a T cell-bilayer interface. Infection status of donors are indicated at the bottom of the plots. Mean values from three independent experiments for each donor group are shown. (**B**) Ability of T cells from HIV- chronic HIV+ and ART-treated HIV+ individuals to form distinct interface structures are compared within CD27+ CD8 T cell subset. The data represents mean values (±SD) of three independent experiments for each donor’s group with different infection status. Statistically significant difference between subject group was determined using one-way ANOVA with Tukey’s post-test; *p<0.05, **p<0.01 and ***p<0.001.

### Purified naïve CD8 T cells from HIV-infected patients exhibit enhanced capacity to form mature synapses

Because CD27+ CD8 T cells contains naïve and more differentiated T cells, it is essential to determine whether purified naïve T cells from patients infected with HIV would still have a greater capacity to form mature synapses as compared to naïve T cells derived from uninfected donors. To this end, we purify naïve T cells as well as transient memory T cells (TM) and effector memory (EM) T cells from HIV-infected people **(Supplemental Figure 8)** and analyzed the formation of synaptic interface by each category of these T cells. As evident from **Figure 5A**, naïve T cells from HIV+ patients demonstrated enhanced capacity to form mature synapses as opposed to those T cells derived from uninfected donors. The fractions of T cells that form mature synapses by TM and EM T cells were similar.

**Figure 5.**
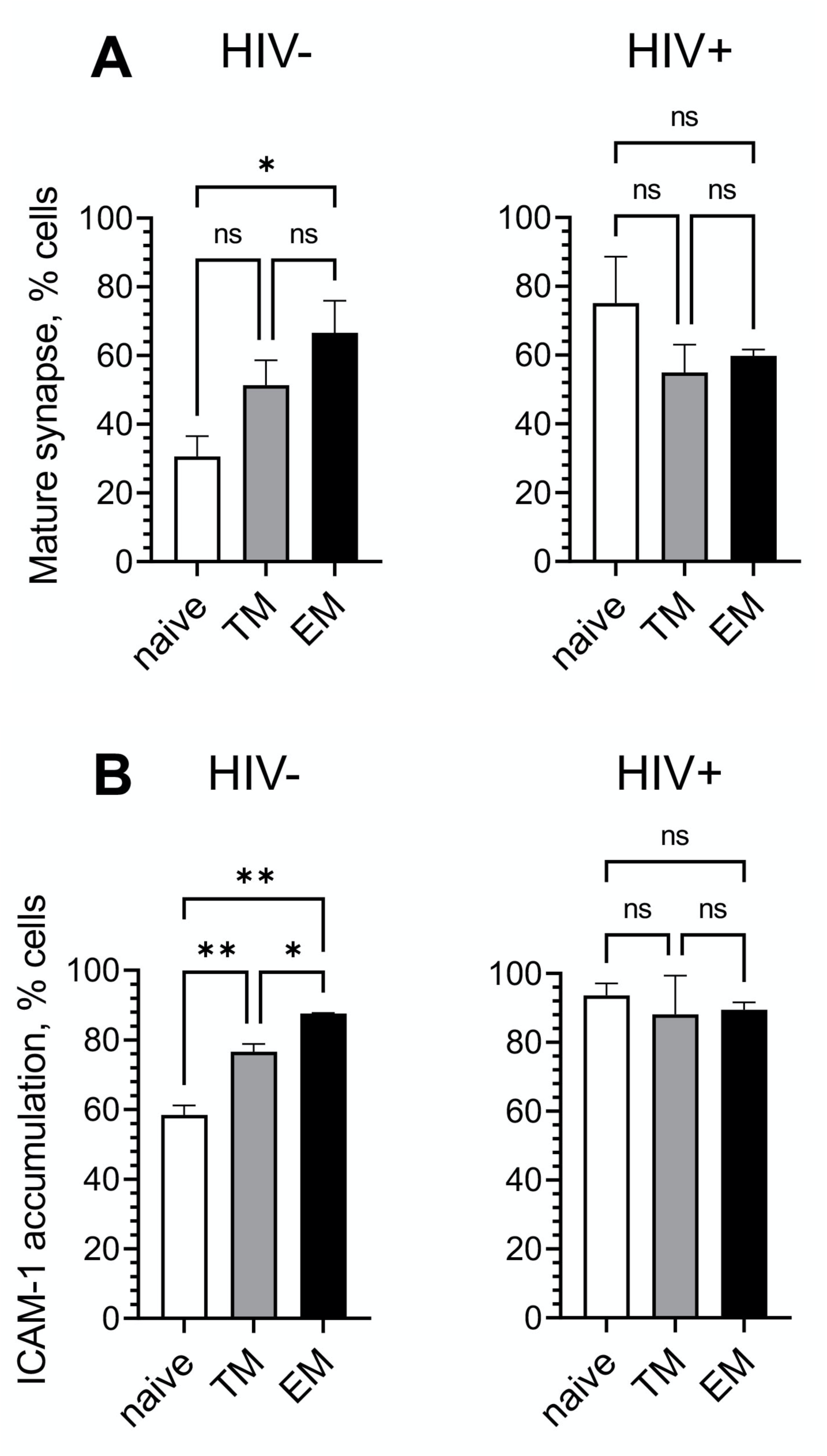
Naive CD8 T cells from chronic HIV+ individuals exhibit greater capacity to accumulate ICAM-1 and form mature synapse compare to T cells from HIV- individuals. Naïve, transitional and effector memory CD8 T cells was sorted out from PBMC of HIV- and chronic HIV+ individuals by flow cytometry as described in **Supplemental Figure 8**. The cells were exposed to bilayers containing anti-CD3 antibodies and ICAM-1, and formation of interface structures were observed by confocal microscopy for 30 minutes. Frequency of naïve, transitional and effector memory CD8 T cells establishing mature synapse (**A**) and accumulating ICAM-1 molecules at synaptic interface (**B**) are shown. Infection status of individuals are indicated. For each condition, bar represent standard deviation of the mean of at least two independent experiments; *p<0.05, **p<0.01 as established by one-way ANOVA with Tukey’s post-test.

Consistent with these results, naïve T cells from the infected individuals revealed increased ability to accumulate ICAM-1 at the T cell/bilayer interface as opposed to naïve T cells from uninfected donors (**Fig. 5B**). The extent of ICAM-1 accumulation by TM and EM T cells was very similar.

These data demonstrate that naïve T cells from HIV-infected individuals revealed behavior of activated T cells.

### Degranulation pattern of CD8 T cell is linked to the type of synaptic interface

We analyzed degranulation patterns of CD8 T cells that establish various types of synaptic interface (**Fig. 6**). Granule release in mature synapses was mostly observed in the cSMAC zone as previously described (**Fig. 6A** and **movie 5**) ^10^. The amount of released granules is steadily increased over time in the cSMAC area (**Fig**. **6E**). T cells establishing focal synapses demonstrated a weak degranulation observed in a small area around CD3/TCR accumulation (**Fig 6B** and **movie 6**). Granule release is linked to weak and transient ICAM-1 accumulation and decreased with time (**Fig. 6F**). Release of cytolytic granules in T cells demonstrating kinapse formation was located in the area of initial attachment of the cell to the bilayer that is left behind and continued in the leading edge of moving cells (**Fig. 6C,G** and **movie 7**). T cells that form multifocal synapses revealed dispersed pattern of granule release (**Fig. 6D and H,** also see **movie 8**). These patterns were seen in T cells at all differentiation stages and independent on donors’ status but were observed at different proportions.

**Figure 6.**
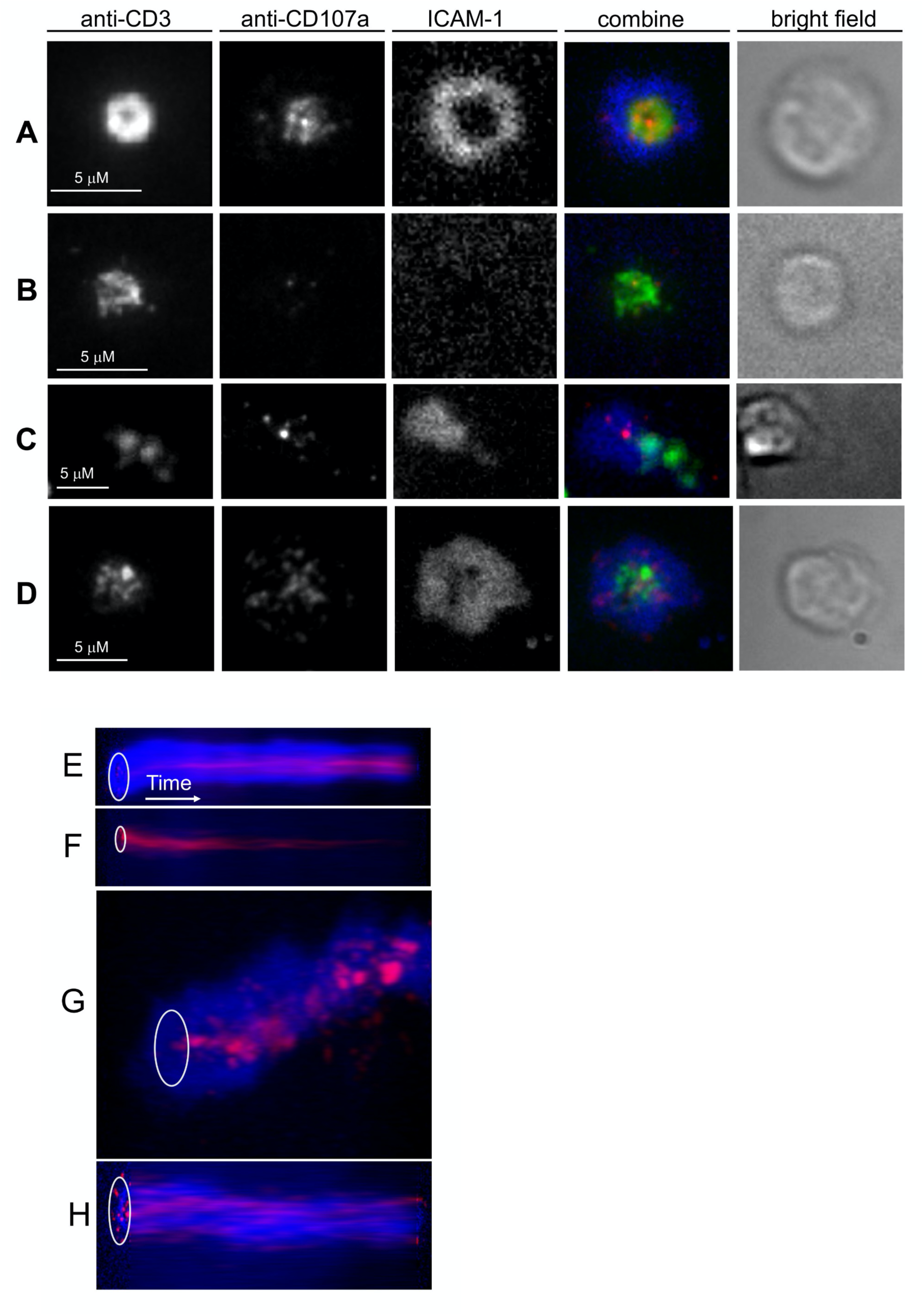
Structure of CD8 T cell-bilayer interface is linked to patterns and kinetics of T cells degranulation. CD8 T cells were purified by negative magnetic sorting and exposed to the bilayers containing anti-CD3 antibodies and ICAM-1 in the presence of anti-CD107a Fab fragments. Structure of the interface and cellular degranulation was observed for half an hour after the initial cell contacts with the bilayers using TIRF (anti-CD3 and anti-CD107a antibodies) and wide field fluorescence (ICAM-1) microscopy. (**A-D**) Representative images show relative positioning of degranulation foci (red), andi-CD3 antibodies (green) and ICAM-1 molecules (blue) at the interface of T cells that form (**A**) mature synapse, (**B**) TCR/CD3 focal synapse, (**C)** kinapse and (**D)** multifocal synapse. Images were taken at 30 minutes after T cell loading on bilayers. ICAM-1 accumulation was visualized by wide field fluorescent microscopy; anti-CD3 antibody clustering and granule release locations were imaged by TIRF microscopy. Scale bars designate 5 μm. (**E-H**) Representative kymograms show temporal changes in spatial positioning of ICAM-1 (blue) and granules (red) released at interface of CD8 T cells forming mature synapse (**E**), TCR/CD3 focal synapse (**F**), kinapse (**G**), and multifocal synapse (**H**). Images were taken at the rate of one frame/min for 30 minutes. White ovals indicate position of cell-bilayer interfaces, white arrows show time axis. Real-time degranulation dynamics shown in **Movies 5-8**.

### Naïve CD8+ T cells from HIV-infected patients revealed atypical granule release

Naive T cells from the infected people were capable to form a higher number of mature synapses that are linked to larger amount of released granules as compared to TCR/CD3 focal synaptic interface (**Fig. 7A**). In addition, the released granules within mature synapses are focused (**Fig. 7B**). Therefore, naïve T cells from HIV infected peoples revealed higher functional activity. In less differentiated T cells (naïve or TM phenotype) total fluorescent intensity of CD107a staining was significantly lower as oppose to cells at later differentiated stages (**Fig 7C**). This was independent of donor status. T cells at later differentiation stages developed in the infected people demonstrated higher capacity to form multifocal synapse leading to a scattered pattern of CD107a staining (**Supplemental Figure 9**). No difference was observed between mature synapses and multifocal synapse in the amount of released granules but area of degranulation within multifocal synaptic interface was significantly higher for T cells demonstrating dispersed degranulation pattern. In contrast, mature synapses revealed focus degranulation pattern in smaller area that reflects more potent functional activity (see **Supplemental Figure 9A** and **B**).

**Figure 7.**
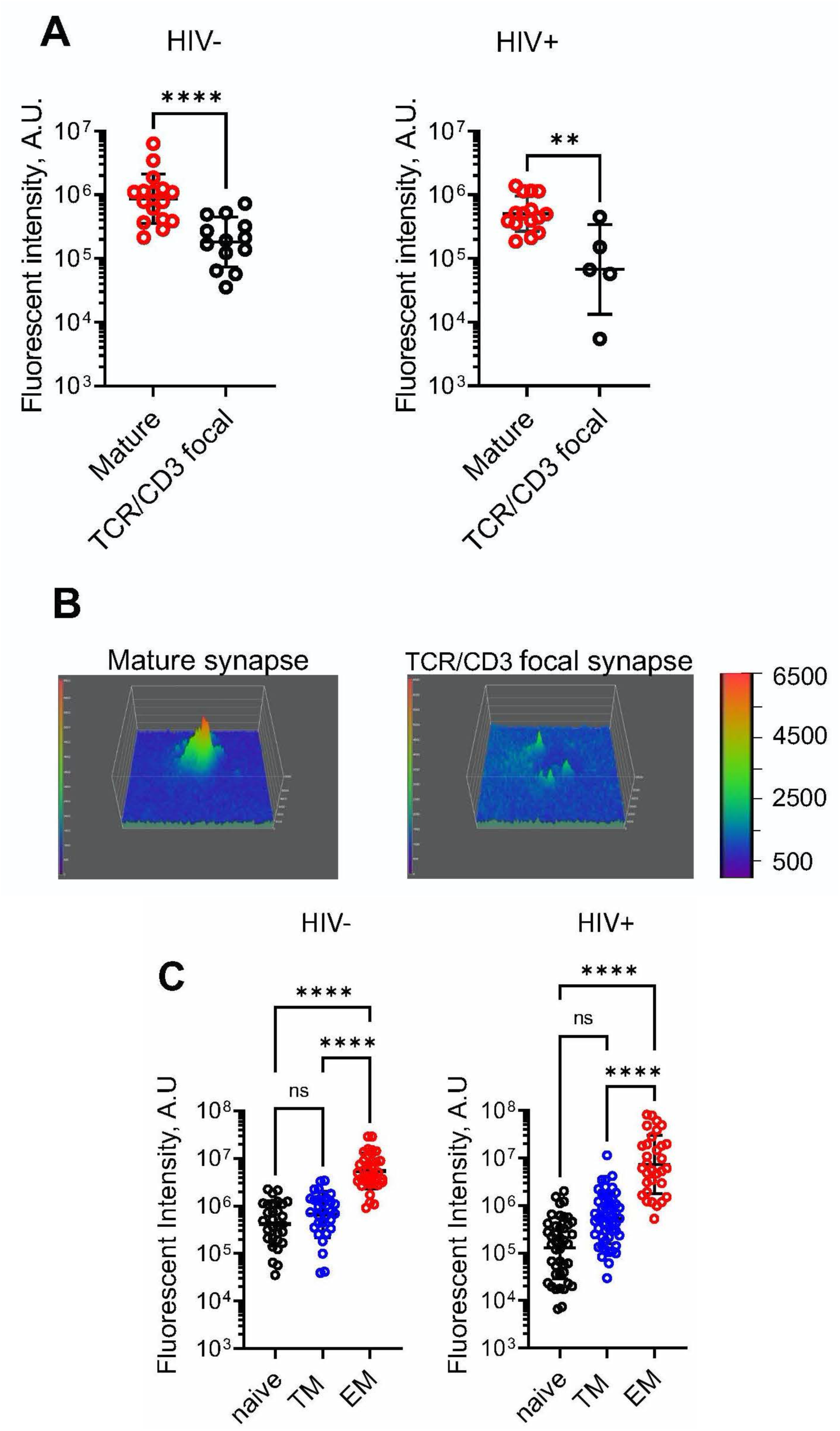
Intensity and quality of degranulation depends on stage of T cell differentiation and type of synaptic interface. Naïve, TM and EM CD8 T cell subsets were isolated by flow cytometry and placed on bilayers containing anti-CD3 antibodies and ICAM-1 in the presence of anti-CD107a Fab fragments. Granule release location was observed by TIRF microscopy, ICAM-1 accumulation was imaged with wide-field microscopy. Images were taken at 30 minute of initial T cell contact with bilayer surface. (**A**) CD107a levels measured at mature synapses (red empty circles) and TCR/CD3 focal interfaces (black empty circles). Infection status of donors indicated at the top of the plot. Representative results from three (HIV+ donors) and two (HIV- donors) independent experiments are shown. Geometric mean ± SD are indicated by black lines; **p<0.01, ****p<0.0001 were determined by non-parametric two-tailed Mann– Whitney test. (**B**) Representative surface intensity plot of naïve CD8 T cell degranulation at mature synapse (**left**) and TCR/CD3 focal interface (**right**). The plots are shown using pseudo-color scale. (**C**) Quantitation of CD107a accumulation at the interface of naïve (black empty circle), TM (blue empty circle) and EM (red empty circle) CD8 T cells from HIV- (left) and chronic HIV+ (right) individuals. Representative results from one of three independent experiments for each group are shown. Each circle represents individual T cell, n≥32. Geometric means are indicated by black lines, error bars represent SD. ****p<0.0001 as determined with non-parametric two-tailed Mann–Whitney test.

## Discussion

Our major findings described here show that majority of naïve CD8 T cells from chronically HIV-infected people form mature synapses demonstrating behavior of activated T cells despite having naïve phenotype. In marked contrast, most of naive T cells from uninfected individuals form focal synapses demonstrating aggregation of TCR but inability to aggregate and segregate LFA-1 (**Fig. 5A**). This is consistent with observation that propensity of naïve CD8 T cells from healthy donors to form synaptic structures significantly reduced as opposed to memory T cells^22^.

The formation of mature synapses is mediated by TCR-dependent conversion of LFA-1 to high-avidity form following interaction with ICAM-1 that leads to cytoskeleton remodeling, segregation of highly polymerized actin to the periphery of the T cell contact area, the actin clearing from the middle of the interface, and the formation of peripheral ring junction ^13, 20, 23^. The expression of LFA-1 in CD8 T cells depends on stage of differentiation with naïve T cells presenting lowest level of LFA-1 molecules on the cell surface (**Fig 3B**). In blood, LFA-1 on naïve T cells is presented in low affinity form and could be converted to higher affinity conformation after engageing CCR7 receptors with chemokines during entry of the T cells into lymph nodes ^24^. However, signaling through chemokine receptors is not sufficient to induce formation of mature synapses by naïve T cells ^25^ most likely due to inability to form high avidity LFA-1 microclusters. Low affinity form of LFA-1 on naïve T cells persists for the first several hours during T cell priming by antigen-presenting dendritic cells resulting in multiple transient contacts of the T cells with APCs allowing preferential activation of T cells with high affinity TCR for the antigen ^26^.

Similar to the cells from healthy donors, naïve CD8 T cells from HIV-infected donors presented low level of LFA-1 molecules ^27^. Despite the low level of LFA-1, naïve T cells from infected individuals have greater ability to accumulate ICAM-1 and form mature synapses (**Fig. 5**) suggesting that chronic inflammation during HIV infection ^28^ could influence productive LFA-1-mediated cytoskeleton remodeling. It has been shown that formation of mature synapses decrease TCR activation threshold ^23^. Thus, enhanced ability to form mature synapses during first phase of naïve CD8 cells priming may result in stimulation of antigen-specific CD8 T cells with lower affinity.

Ability of naïve CD8 T cells from HIV infected donors to form mature synapses resembles the behavior of memory T cells (**Fig 5**). It has been shown that naïve T cell activation preferentially initiates ERK signaling pathways through SLP76 scaffolding molecule, while in memory T cells TCR stimulation induces alternative p38 pathway. ^29^. Activated ERK acts as negative regulator of LAT/SLP-76 signalosomes facilitating NF-κB signaling and calibrating initial response of naïve CD8 T cells to antigen of different strength ^30, 31^. In contrast, hDlg-dependent phosphorylation of p38 leads to activation of NFAT that is linked to production of cytokines such as IL-2 and INF-γ ^32, 33^. Scaffold protein hDlg is recruited to immunological synapse in response to TCR/CD28 enagagement and facilitates interaction of Lck with ZAP70 and WASp coordinating actin-mediated synapse formation and effector function ^33^. Dlgh1 has been shown to form a dynamic complex with actin-binding protein 4.1G and LFA-1-binding protein CD226 (DNAM-1) that leads to formation of high avidity LFA-1 clusters required for mature synapse formation. Thus, TCR stimulation on naïve T cells from HIV-infected people appears to induce preferentially alternative p38 dependent signaling pathway that links conversion of LFA-1 to high affinity form, formation of mature synapses and degranulation (**Fig. 5**).

Until recently, naïve T cells have been considered as ‘deeply’ quiescent and mostly homogeneous subset ^34^. However, recent data point out that naïve T cells are heterogeneous population, and subset(s) of T cells with ‘classical’ naïve phenotype could demonstrate transcriptional profile of more differentiated T cells ^35, 36^. Such naïve T cells can also produce various cytokines and are more sensitive to antigen stimulation ^36, 37^. Unconventional naïve CD8 T cells often express chemokine receptor CXCR3, which could direct them to inflamed sites. Expression of CXCR3 is associated with increased expression of accessory molecules CD226 that is involved in LFA-1-mediated costimulatory signals for triggering naive T cell differentiation and proliferation ^35, 36, 38–40^. Partial differentiation of naïve cells has been attributed to homeostatic proliferation of peripheral naïve T cells that arises due to thymic output of naïve quiescent T cells with high TREC content. During untreated HIV infection, homeostatic proliferation of naïve CD8 T cells increases three-fold followed parallel increase of CD8 T cell loss ^41^. Effective antiretroviral therapy has a reverse effect but did not completely normalize turnover of naïve T cells ^42^.

Even though the heterogeneity of naïve T cells could be mediated by multiple factors including thymic output, chronic inflammation and aging ^43^, continuous inflammation during HIV infection is a major factor that influence proliferation, graduate loss of stemness properties and partial differentiation of naïve T cells ^44, 45^. We suggest that these conditions promote appearance of high avidity LFA-1 on naive T cells that is mediated by alternative p38-dependent signaling pathway, synapse formation and degranulation. Indeed, the presence of naïve T cells with a higher functional activity in HIV-infected people is evident from their degranulation capacity. While majority of naïve T cells from healthy donors are characterized by faint and transient CD107a accumulation (**Fig. 7B and C** and **movie 6B**), naïve CD8 T cells from HIV- infected individuals revealed sustained and focused CD107a staining which was similar to memory T cells (**Fig. 7C and Supplemental Figure 9B**). However, the observed CD107a staining was less bright than that of effector memory T cells indicative of a different content of released granules (**Fig. 7A)** ^46, 47^.

Chronic inflammation also affects plasticity of APC due to the increase of cellular stiffness ^48, 49^. This enhances their capacity to stimulate T cells changing T-cell metabolic properties and cell cycle progression. At the same time, fluidity of the cell membrane become greater as a result of substantial changes of lipid composition of the membrane ^50^. Membrane fluidity that depends on orderliness of membrane lipids facilitates rearrangement of cell surface proteins and formation of highly ordered synaptic interface. In addition, IFN-γ induces increase of the expression level of MHC proteins and integrin ligands on antigen presenting cells that enhances their stimulatory potency.

Transitional memory (TM) cells from both HIV+ and HIV- donors were capable to form mature synapses and revealed focal degranulation (**Fig. 5** and **7C**). TM cells are tissue trafficking cells that can access lymphoid and non-lymphoid tissues and are thought to be non-cytolytic ^51^. The latter is consistent with less dense CD107a staining at the synaptic interface. Importantly, TM cells in HIV-infected people represent significantly higher fraction in circulation as compared to uninfected donors (Betts et al., unpublished observation) suggesting that TM cells could be readily developed from perverted naïve T cells during HIV infection.

Although ART-treatment of HIV-infected patients significantly suppresses HIV replication, reduced frequency and elevated homeostatic proliferation of naïve CD8 T cells is still observed ^52, 53^. Consistent with this, we noticed enhanced ability to accumulate ICAM-1 at synaptic interface by early differentiated CD8 T cells from ART treated donors similar to that observed for naïve T cells from HIV infected untreated patients (**Fig 2**). Even though ART treatment diminished ability of early differentiated cells to form mature synapses, significant fraction of these cells still form synapses albeit with more transient mode (kinapses) as compared to the cells from donors with uncontrolled HIV (**Fig. 6**). The motile cells have lamellipodium upfront that contain most of redistributed and accumulated ICAM-1 in a central lamella region followed by a trailing uropod concentrating engaged TCRs. Degranulation at kinaptic interface is not focused, and released granules dispersed along the cell trajectories suggesting impairment in granule delivery. It has been shown that formation of immunological kinapses reflects reduced antigen sensing by T cells ^54, 55^. Thus, early differentiated CD8 T cells are functionally less activated, and fraction of naïve T cells that possess more ‘quiescent’ properties is increased followed antiretroviral therapy. How duration of the treatment and age of the patients modulate synaptic interface structure of naïve CD8 T cells is unclear and is a topic of future studies.

At late stages of differentiation, significant proportion of T cells from infected people demonstrated formation of multifocal synapses with scattered pattern of CD107a staining as opposed to the focused staining typical for CD8 T cells capable to mount efficient cytolytic response (**Supplemental Figure 9**) ^10, 11^. The structure is characterized by well-developed highly dynamic lamellipodium that persists during observation period in contrast to the formation of mature synaptic interface where lamellipodium quickly disappeared following initial T cell spreading. The dispersed degranulation pattern was often observed at synaptic interface formed by NK cells ^56, 57^. At these conditions, cytolytic granules are released over the synaptic interface at multiple locations devoid of highly polymerized actin ^58^. We have previously shown that formation of stable mature synapses with focused granule delivery increase the efficacy of target cell killing by cytotoxic CD8 T cells in several folds ^10, 23^. In contrast, we would like to suggest that terminally differentiated CD8 T cells that form highly motile synapses with scattered granule release have diminished ability to exercise effector functions. These T cells may represent senescent effector CD8 T cells that are developed in chronically infected HIV patients ^59, 60^. Increased accumulation of dysfunctional CD8+ T cells that reveal phenotype of senescent cells including cell cycle arrest, short telomeres, and diminished of CD28 surface expression is linked to severity of HIV disease. The abundance of CD8+CD28-T cells early in the infection is predictive of progression to AIDS ^61^.

Increase of senescent cell population together with reduction in naïve CD8 T cells is hallmark of aging of the immune system. It is likely that the CD8+ T cell defects caused by HIV infection may synergize with similar defects associated with aging further solidifying needs to apply latest techniques for more thorough characterization of these T cells subsets ^62, 63^.

## METHODS

### Human blood samples

Peripheral blood samples were collected from HIV negative donors and individuals with chronic HIV infection untreated or treated with ART. All participants enrolled in this study provided written informed consent, which was approved by the regional ethical research boards and conform to Helsinki Declaration. Subject clinical parameters are summarized in S1 Table. The interval of donors’ age in each subgroup was relatively narrow (range 19-39 years).

### Cells

Hybridoma OKT3 producing antibodies recognizing human CD3ε and hybridoma TS2/4.1.1 secreting antibodies against human LFA-1 were purchased from ATCC. Hybridoma H4A3 secreting antibody against CD107a (LAMP-1) was kindly provided by Dr. J. Thomas August, Department of Pharmacology and Molecular Sciences, Johns Hopkins Medical School. Peripheral blood mononuclear cells (PBMC) were isolated from whole blood using Ficoll-Hypaque density gradient centrifugation and cryopreserved at −140°C. Hybridoma YN1.1 producing antibodies against ICAM-1 was kind gift from Michael Dustin, Skirball Institute of Biomolecular Medicine, New York University.

### Antibodies and proteins

Antibody against CD107a was purified from hybridoma H4A3 supernatant. Monovalent Fab fragments of CD107a-specific antibody were produced by papain digestion, purified by anion exchange chromatography and labeled with Alexa Fluor 568 as previously described (Beal et al., 2008; Steblyanko et al., 2018). Anti-CD3 antibodies (clone OKT3) were purified from cell culture supernatant. The antibody was mono-biotinylated and labeled with Alexa Fluor 488 as described (Steblyanko et al., 2018). Mouse anti-human antibody against LFA-1 were purified using hybridoma TS2/4.2.1.1 and labeled with Alexa Fluor 488.

Recombinant soluble ICAM-1 protein was expressed in a *Drosophila Melanogaster* cells using pMT/V5-His vector with inducible promotor (Invitrogen), purified by sequential two-step affinity chromatography on anti-ICAM-1 antibody (clone YN1.1) Sepharose and Ni-NTA agarose, and labeled with Cy-5 dye as previously described (Steblyanko et al., 2018). The affinity Sepharose resin was prepared by coupling purified rat anti-mouse YN1.1 antibody with CNBr-activated Sepharose 4B (Sigma).

The following mouse anti-human antibodies were used for purification of CD8 T cell subsets by Flow Cytometry: anti-CD45RO PE CF594 (clone UCHL1, BD Biosciences), anti-CD56 PE Cy7 (clone B159, BD Biosciences), anti-CD45RA BV650 (clone HI100; BD Biosciences), anti-CCR7 APC Cy7 (clone G043H7, Biolegend), anti-CD27 BV785 (clone O323, Biolegend), anti-CD10 BV605 (clone HI10A, Biolegend), anti-CD14 BV510 (clone M5E2, Biolegend), anti-CD19 PE (clone HIB19, Biolegend), anti-CD16 BV510 (clone 3G8, Biolegend) and anti-CD4 PE Cy7 (clone RPA-T4, Biolegend). The LIVE/DEAD Fixable Aqua Dead Cell Stain Kit (Invitrogen) was used for exclusion of dead cells. Anti-CD3 and anti-CD8 antibodies were excluded from the panel to avoid CD8 T cell stimulation during the staining. After magnetic sorting, purity of CD8 T cells and their subsets were confirmed by Flow cytometry analysis with anti-CD8, anti-CD45RO and anti-CD27 antibodies.

### Isolation of CD8 T cells and CD8 T cell subsets

Cryopreserved PBMC were thawed and rested overnight at 2 x 10^6^ cells/ml in R10 medium (RPMI with 10% fetal bovine serum, 1% penicillin-streptomycin and 1% L-glutamine) with 10 units/ml of DNAse I (Sigma-Aldrich) at 37°C and 5% CO_2_. After resting, cells were washed in PBS and used for isolation of CD8 T cells or CD8 T cell subsets by magnetic or flow cytometry sorting.

#### Magnetic sorting

CD8 T cells were purified from PBMC by negative immunomagentic sorting using CD8+ T Cell Isolation Kit (Miltenyi Biotec) and used for bilayers experiments or for further separation into the subsets. CD27+ CD8 T cells were separated with CD27 MicroBeads (Miltenyi Biotec). Negatively selected population were incubated with CD45RO MicroBeads for second round of separation on CD27-CD45RO+ and CD27-CD45RO-CD8 T cell subsets. The cells were washed twice in the assay buffer (20 mM HEPES, pH 7.4, 137 mM NaCl, 2 mM Na_2_HPO4, 5 mM D-glucose, 5 mM KCl, 2 mM MgCl_2_, 1 mM CaCl_2_, and 1% human serum albumin). After counting the cells were resuspended in the assay buffer at density 2x10^6^/ml and kept on ice prior to experiments. The purity of the subsets (>90%) were confirmed by flow cytometry analysis using anti-CD8, anti-CD45RO and anti-CD27 antibodies.

#### Flow cytometry sorting

PBMC were pre-stained with antibody against CCR7 in PBS buffer for 10 minutes at 37°C. All following incubations were performed at room temperature in the dark. Cells were stained for viability exclusion with LIVE/DEAD Aqua for 10 minutes. Optimized antibody cocktail was prepared in FACS buffer (0.1% sodium azide and 1% bovine serum albumin in 1X PBS) and was combined with washed cells for 20 minutes to detect additional surface markers. The cocktail contained mouse anti-human antibodies against CD45RO, CD56, CD45RA, CD10, CCR7, CD27, CD14, CD19, CD16 and CD4. Cells were washed with FACS buffer and resuspended in 350 µl of phenol red-free RPMI. Cells were sorted using a FACS ARIA II (BD Biosciences) with low pressure settings into 1.5 mL DNA LoBind tubes (Eppendorf) pre-filled with 300 μl of R10. The gating strategy is shown in the **Supplemental Figure 8**.

### Measuring LFA-1 expression level

CD8 T cells or purified CD8 T cell subsets were stained with anti-LFA-1 antibody labeled with Alexa Fluor 488 at concentration 2 μg/ml, and the fluorescent intensity were analyzed by Flow Cytometry. Fluorescent intensity of Alexa Fluor 488 calibration beads with defined number of the fluorescent molecules per bead (Bangs labs Quantum MESF kits) was recorded at the same day and instrument settings. The beads MFI was analyzed with FlowJo software and plotted against the numbers of the fluorophore per beads using Bangs Laboratory software. The dependence was utilized for conversion of the cell fluorescent intensity values to numbers of the receptors per cell considering that the Alexa Fluor 488/antibody ratio was 4:1 and one antibody molecule could interact with two receptors.

### Planar lipid bilayers

Planar lipid bilayers were prepared as previously described (Steblyanko et al., 2018; Buggert, et al., 2018). Briefly, liposome mixture containing NiNTA-DGS (Avanti Polar) and biotinyl-CAP-phosphoethanolamine lipids at final molar concentration 17.5% and 0.01% correspondingly was used to form bilayer surfaces on glass surface of sticky-Slide VI 0.4 (ibidi) channels. Streptavidin and biotinylated anti-CD3 antibody labeled with Alexa Fluor 488 were sequentially loaded onto bilayers to produce the antibody density of 50 molecules/μm^2^. Cy5-labeled ICAM-1 molecules were introduced into the bilayers through interaction with Ni-NTA-lipids at final density 300 molecules/μm^2^. Densities of Cy5-ICAM-1 and anti-CD16 antibody on the bilayers were determined as described elsewhere (Anikeeva et al., 2005). Channels of ibidi chamber containing bilayers were washed and kept in the assay buffer (20 mM HEPES, pH 7.4, 137 mM NaCl, 2 mM Na_2_HPO4, 5 mM D-glucose, 5 mM KCl, 2 mM MgCl_2_, 1 mM CaCl_2_, and 1% human serum albumin) prior to use.

### Confocal Imaging

Confocal fluorescent microscopy was carried out on scanning laser Nikon A1R+ confocal microscope with heated stage, objective heater and a motorized stage with autofocus. The ibidi chamber with preformed bilayers was placed on preheated stage and 60X/1.4 oil objective of the confocal microscope. The cell samples were injected into the entry ports of the slide containing bilayer with displayed ligands. The stage positions were chosen during first two minutes after the sample loading. The selected stage positions were imaged every 2 minutes for 30 minutes. Bright-field, reflected light, and two fluorescent channels (Alexa Fluor 488 for anti-CD3 imaging and Cy5 for ICAM-1 imaging) of the confocal microscope were utilized to acquire images of interface formed by CD8 T cells interacting with the bilayer surface.

### TIRF Imaging

TIRF imaging were utilized to assess CD8 T cell degranulation over time. The imaging was performed by Andor Revolution XD system equipped with Nikon TIRF-E illuminator, 100/1.49 NA objective, Andor iXon X3 EM-CCD camera, objective heater, and a piezoelectric motorized stage with Perfect Focus. The objective and stage of the microscope were preheated before placement of the chamber with the bilayers on the stage. Alexa Fluor 568 labeled anti-CD107a antibody Fab fragments were added to the cell suspension at a final concentration of 2 µg/mL, and the cells were injected into the entry ports of an ibidi chamber channel. The stage positions were chosen during first four minutes after cells injection, and the images of the cell-bilayer interface were recorded for 30 minutes at a rate of one image per minute. TIRF mode were used to assess Alexa Fluor 488 (anti-CD3) and Alexa Fluor 568 (anti-CD107a) fluorescence, widefield was utilized to image Cy5 fluorescence (ICAM-1). The cells morphology was assessed with DIC transmitted light.

### Image processing and analysis

To ensure equitable comparison, the measurements of CD8 T cell subsets from individual donors were performed in a series of parallel experiments in the same day using the same reagents and freshly prepared planar bilayers. The images were acquired using the identical microscope settings. Sequential images collected each channel were composed into stacks using MetaMorph image processing software. To determine the parameters of cell-bilayer interactions, we chose only CD8 T cells productively interacting with the bilayers. That was determined by accumulation of anti-CD3 antibodies and formation of adhesion area at the interface and confirmed by morphology analysis of the cells observed in the transmitted light images. Clustered cells and visibly damaged or apoptotic cells were excluded from analysis. Cell was discerned to accumulate ICAM-1, if an increase of Cy-5 fluorescence was observed at least on 10 consequent fields. The extent of ICAM-1 accumulation for selected cells was measured by determining the average fluorescence intensity of accumulated Cy5-labeled ICAM-1 molecules at the cell-bilayer interface over background fluorescence outside of the contact area but in close proximity to the cell. If accumulated ICAM-1 molecules formed a ring structure that was observed on at least two consecutive images, we determined that the cell developed pSMAC. The adhesion area corresponds to tight contact between the cells and bilayer and was observed on IRM images as dark region at interface between cells and bilayers. Regions were drawn around tight adhesion areas on the IRM images, and size of those regions were determined.

Productively interacting cells were divided into four categories in accordance with their characteristics of the interface and morphological features. The cell that formed ‘classical’ pSMAC and cSMAC structures on at least 10 consequent field belongs to mature synapse category. These cells established middle sized adhesion area at the interface and do not form lamellipodia for at least 15 minutes of observation period. CD8 T cells with sporadic or without ICAM-1 accumulation at bilayer surface classified as cells with TCR/CD3 focal interface. These cells form small adhesion area contacts with the bilayers presumably through uropod-like structure that visible on bright field images in some cases. Motile cells forming asymmetric migratory shape with lamella containing accumulated ICAM-1 and uropod with anti-CD3 antibodies accumulation were defined as cells with kinaptic interface. These cells also characterized by a large asymmetrical adhesion area. The symmetrical cells with large adhesion area and visible lamellipodium structure that observed during almost all period of observation belongs to another category defined as multifocal synapse. Multifocal synapse characterized by multiple small accumulations of anti-CD3 antibodies that can partially coalesce into large central area during period of observation.

### DATA AVAILABILITY

All Data will be available after publication of the manuscript and will be limited to non-commercial uses only

## ACKNOWLEDGEMENT

This work was supported by NIH R01 grant by R01AI11869. We are grateful the Sidney Kimmel Cancer Center Bioimaging Shared Resource for excellent support. We also thank the Flow Cytometry Facility of the Sidney Kimmel Cancer Center for excellent technical assistance. We would like to acknowledge help and support of all members of Yuri Sykulev and Mike Betts laboratories.

## AUTHOR CONTRIBUTION

N.A., M.S., M.R.B. and Y.S. conceived of the project, designed, and perform the experiments. N.A., M.S., L.K.-C., M.B. derived and characterize peripheral blood CD8 T cells from HIV- infected and uninfected people. N.A., M.S., M.R.B. and Y.S. analyzed the data and wrote the manuscript

## SUPPLEMENTAL INFORMATION

**Table S1.** Clinical characteristics of HIV-infected donors^*)^

|  | HIV+ | HIV+ ART |
| --- | --- | --- |
| Number of individuals | 8 | 5 |
| CD4 count (cells/ml)<sup>**)</sup> | 382<br>(320-480) | 550<br>(396-720) |
| Viral load<br>(copies/ml)<sup>**)</sup> | 43,789<br>(26,566-43,789) | <50 |

<sup>\*)</sup>The interval of donors' age in each subgroup was relatively narrow (range 19-39 years).
<sup>\*\*)</sup> Median and interquartile range

**Supplemental Figure 1.**
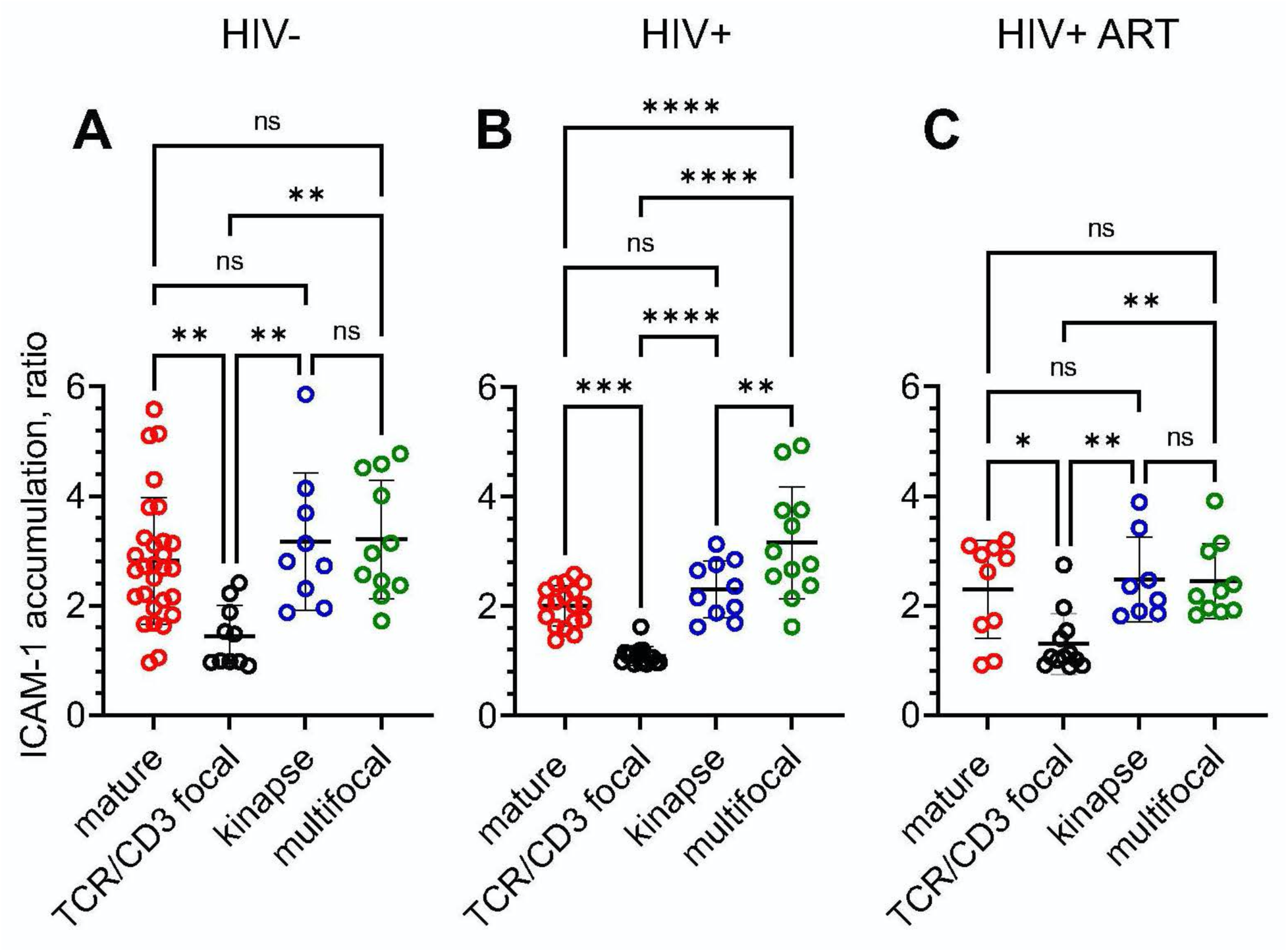
ICAM-1 accumulation is similar at all kind of synaptic interfaces except for TCR/CD3 focal one. CD8 T cells were purified from PBMC of HIV- (**left**), chronic HIV+ (**middle)** and ART-treated donors (**right**) by negative magnetic sorting, and were loaded onto bilayer surface presenting fluorescent-labelled anti-CD3 antibodies and ICAM-1 molecules. The contact interfaces were imaged by confocal microscopy at 10 minutes after the T cell loading. The extent of ICAM-1 accumulation was calculated as a ratio of fluorescent intensity of the ICAM-1 enriched T cell contact area over ICAM-1 fluorescent intensity of a region adjacent to the contact area. The fold difference was calculated for the cells establishing mature synapse (red empty circles), TCR/CD3 focal interfaces (black empty circles), kinapses (blue empty circles) and multifocal synapses (green empty circles). Representative results from one of three experiments for each donor group are shown. Mean ± SD are indicated by black lines and error bars; *p<0.05, **p<0.01, ***p<0.001, ****p<0.0001 as established by one-way ANOVA using Tukey post hoc test.

**Supplemental Figure 2.**
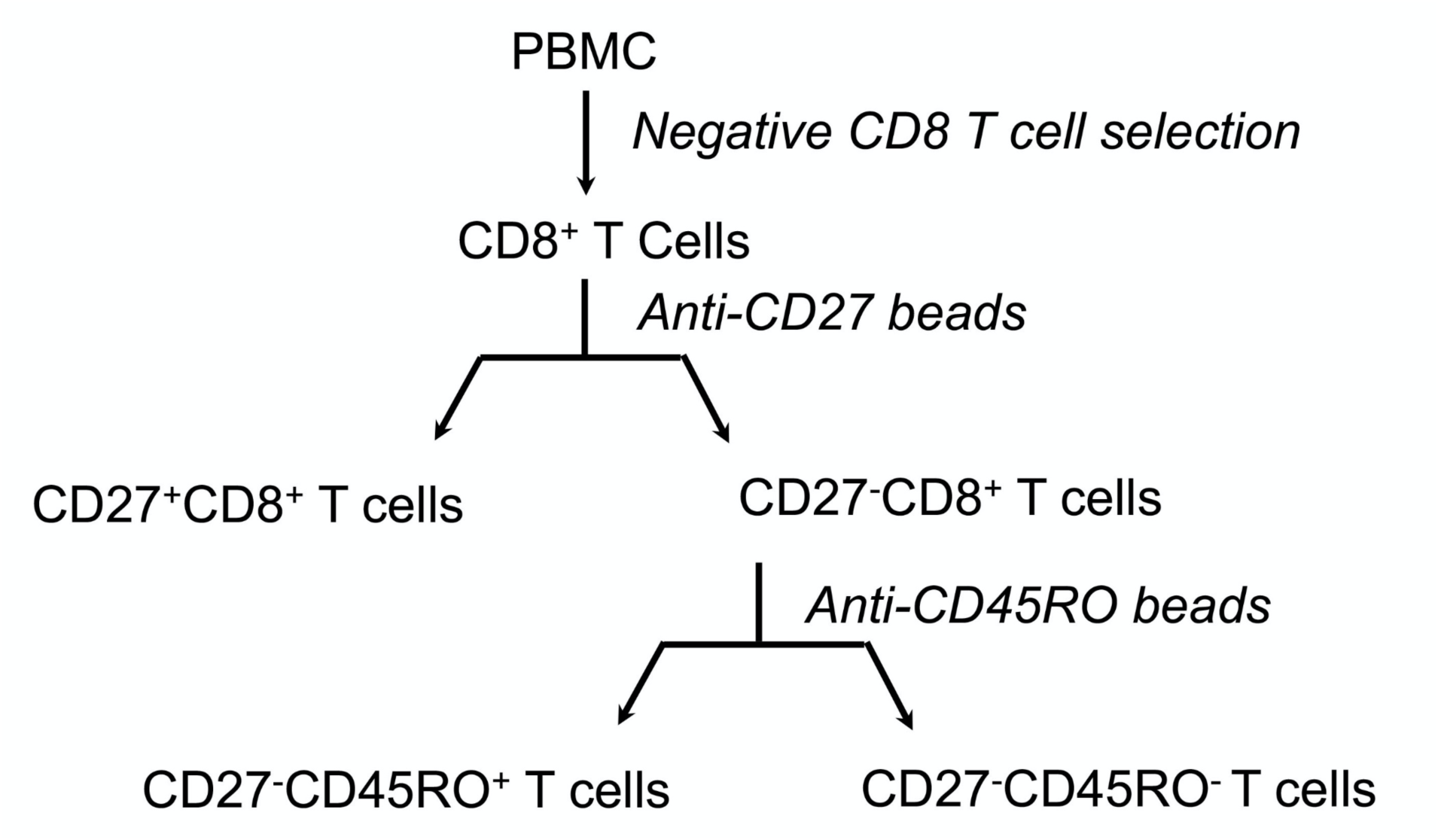
Isolation of CD8 T cells at different stages of differentiation using magnetic sorting. CD8 T cells were isolated from PBMC by negative magnetic sorting. The cells were labelled with anti-CD27 magnetic microbeads and were positively selected using MACS separating column. The eluted fraction represents early differentiated CD27+ CD8 T cells. The CD27- depleted fraction were further labeled with anti-CD45 magnetic microbeads. The second round of separation generated CD27-CD45RO+ CD8 T cell subset and CD27- CD45RO- CD8 T cells at late differentiation stage.

**Supplemental Figure 3.**
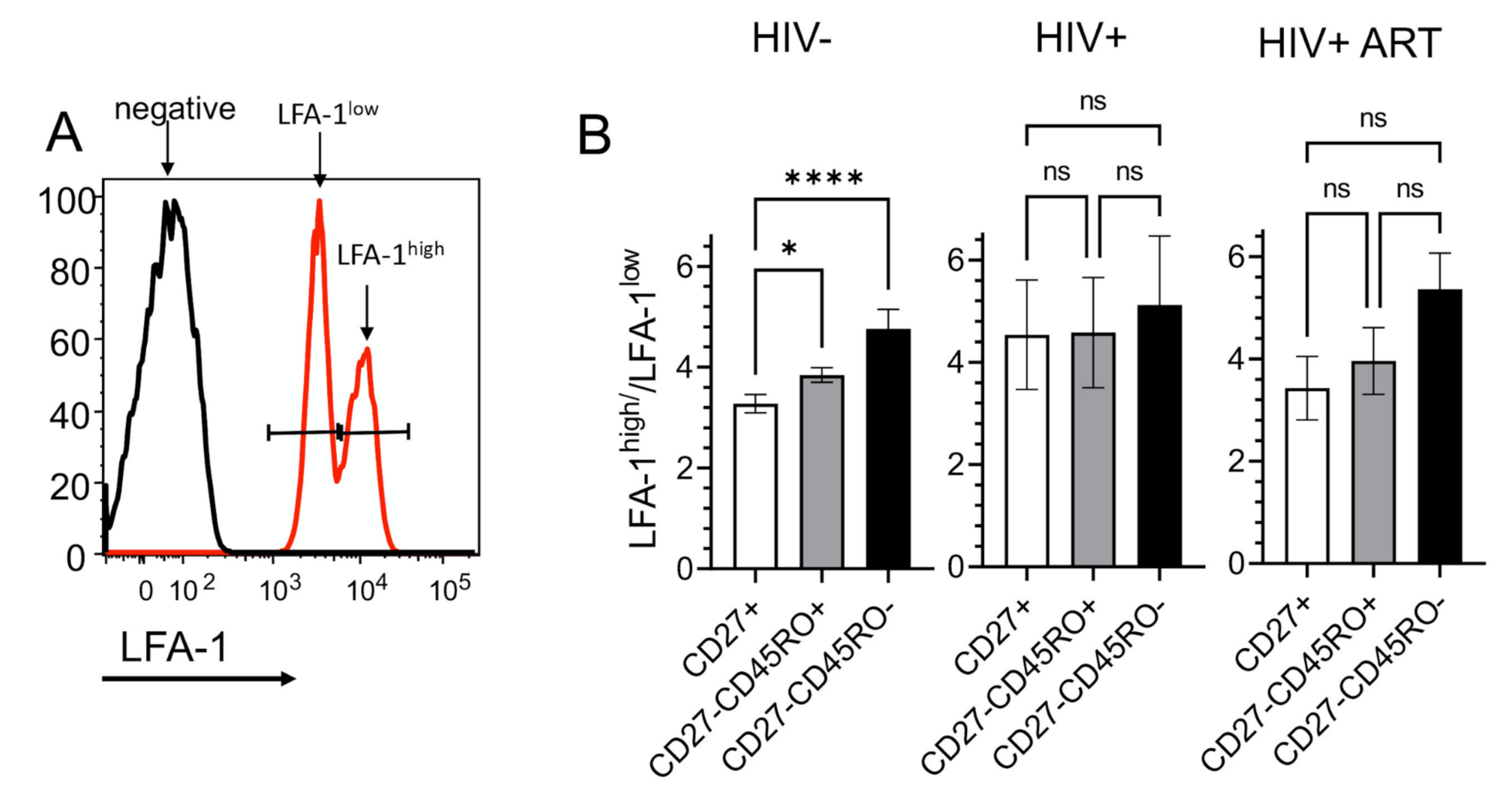
Gating strategy for analysis of T cell fractions with distinct expression level of LFA-1. CD27+, CD27-CD45RO+ and CD27-CD45RO-CD8 T cell subsets were purified from PBMC by magnetic sorting. The cells were stained with 2 μg/ml Alexa Fluor 488 labelled non-stimulatory antibodies against LFA-1 (clone TS2/4) and analyzed by Flow Cytometry. Fluorescent intensity of Alexa Fluor 488 calibration beads (Bangs laboratory) was recorded at the same day and the initial instrument settings was used for quantifying number of LFA-1 molecules per cell. (**A**) Representative histogram shows gating strategy for LFA-1^low^ and LFA-1^high^ fractions CD27+ CD8 T cell subset isolated from HIV- individual. (**B**) Ratio LFA-1^high^/LFA-1^low^ molecules per cell for CD8 T cells at various differentiation stages derived from HIV-, HIV+, and ART-treated HIV+ people. The same gate as shown in panel (**A**) were used for the analysis of the T cells.

**Supplemental Figure 4.**
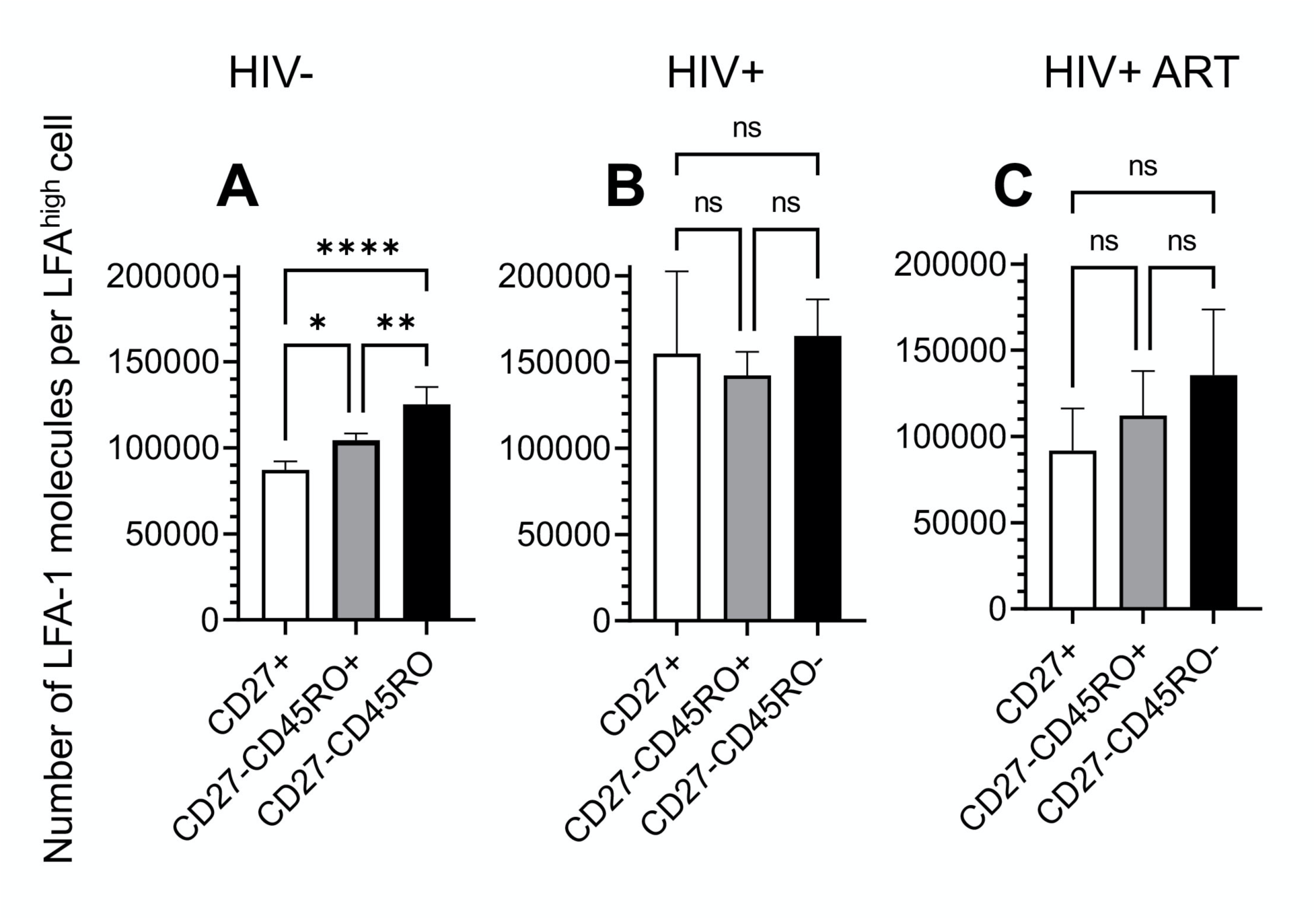
Quantitation of LFA-1 expression level on CD8 T cell subsets. CD27+, CD27-CD45RO+, and CD27-CD45RO-CD8 T cell subsets from HIV- (**A**), HIV+(**B**), and ART-treated HIV+ (**C**) individuals were analyzed. The T cells were stained with either anti-LFA-1 (clone TS2/4) or isotype matched antibodies labeled with Alexa Fluor 488. Fluorescent intensity of Alexa Fluor 488 calibration beads (Bangs laboratory) were exploited at the same instrument settings to quantify number of LFA-1 molecules per cell. For each condition, bar represent standard deviation of mean of three independent experiments; *p<0.05, **p<0.01 and ****p<0.0001 as established by one-way ANOVA with Tukey post hoc test.

**Supplemental Figure 5.**
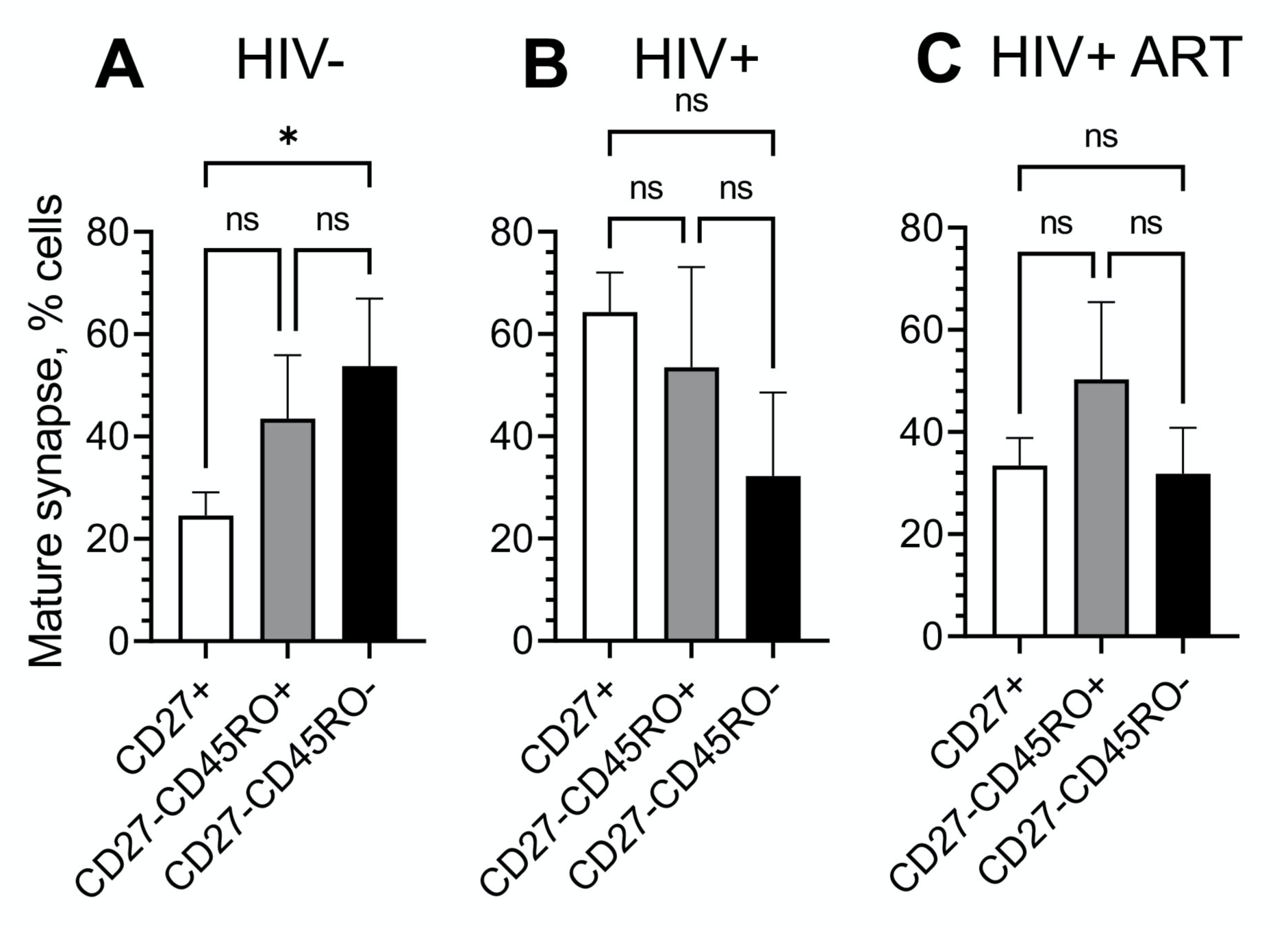
Ability to form mature synapse during differentiation have upward trend only for CD8 T cells from HIV- donors. CD8 T cell subsets from HIV- (**left**), HIV+(**middle**), and ART-treated HIV+ (**right**) individuals were exposed to bilayer surfaces presenting anti-CD3 antibodies and ICAM-1 molecules. Serial images of T cell/bilayer interface were taken for 30 min by confocal microscopy, and number of the cells establishing mature synapses were determined. The results from at least three independent experiments for each donor group are presented as mean ± SD; *p<0.05 by one one-way ANOVA with Tukey post hoc test.

**Supplemental Figure 6.**
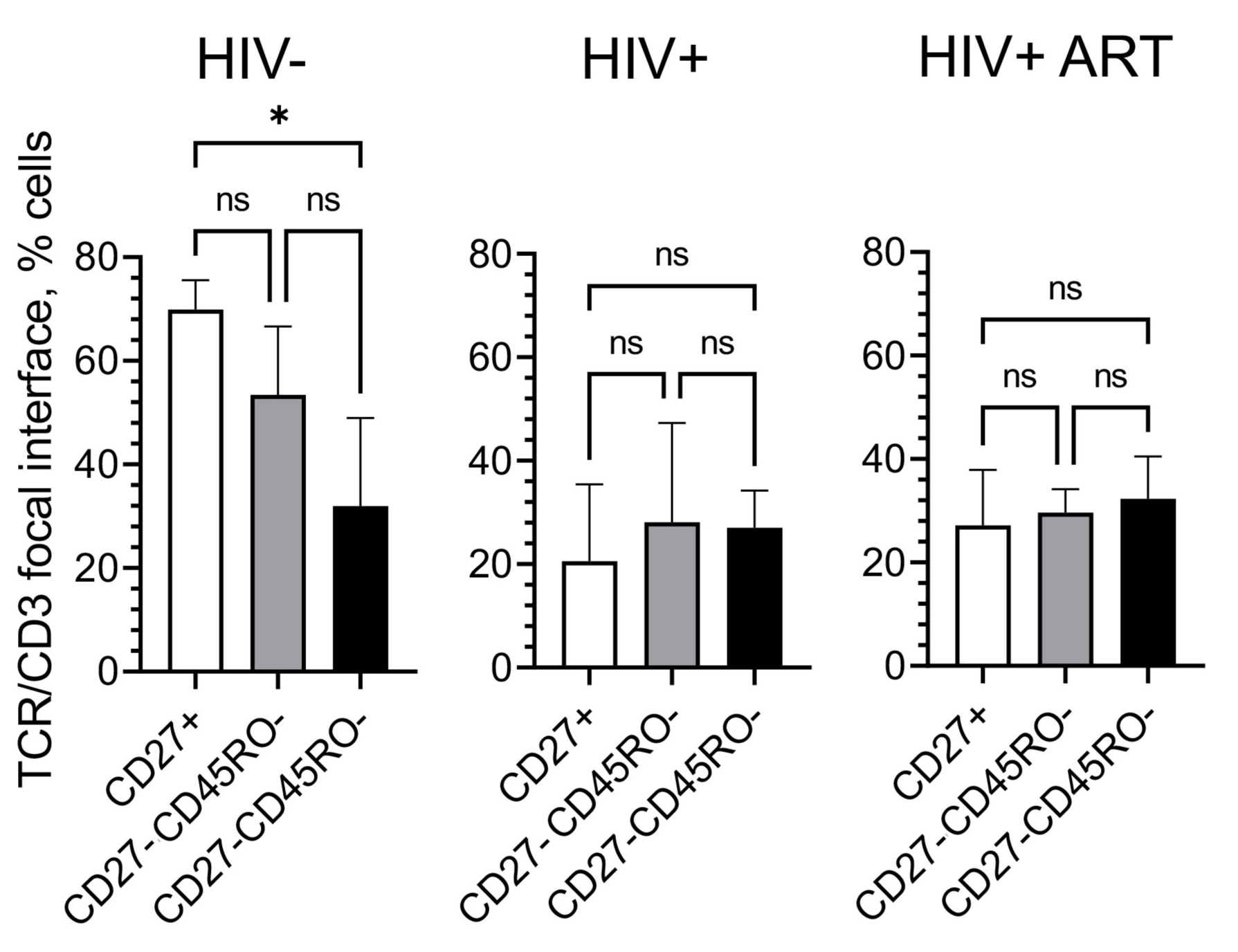
In contrast to CD8 T cells from HIV- donors, the T cells from ART-treated and untreated HIV+ individuals revealed similar fractions of T cells with CD3/TCR focal interfaces regardless of differentiation stage. CD8 T cell subsets from HIV- (**left**), HIV+(**middle**), and ART-treated HIV+ (**right**) individuals were exposed to bilayer surfaces presenting fluorescent-labeled anti-CD3 antibodies and ICAM-1 molecules. The formation of synaptic interfaces was observed for 30 minutes by confocal microscopy at rate 0.5 frame/min. Number of the cells capable to form CD3/TCR focal interface were determined. At least three independent experiments were done for each donor group. Mean ± SD are indicated; *p<0.05 by one one-way ANOVA with Tukey post hoc test.

**Supplemental Figure 7.**
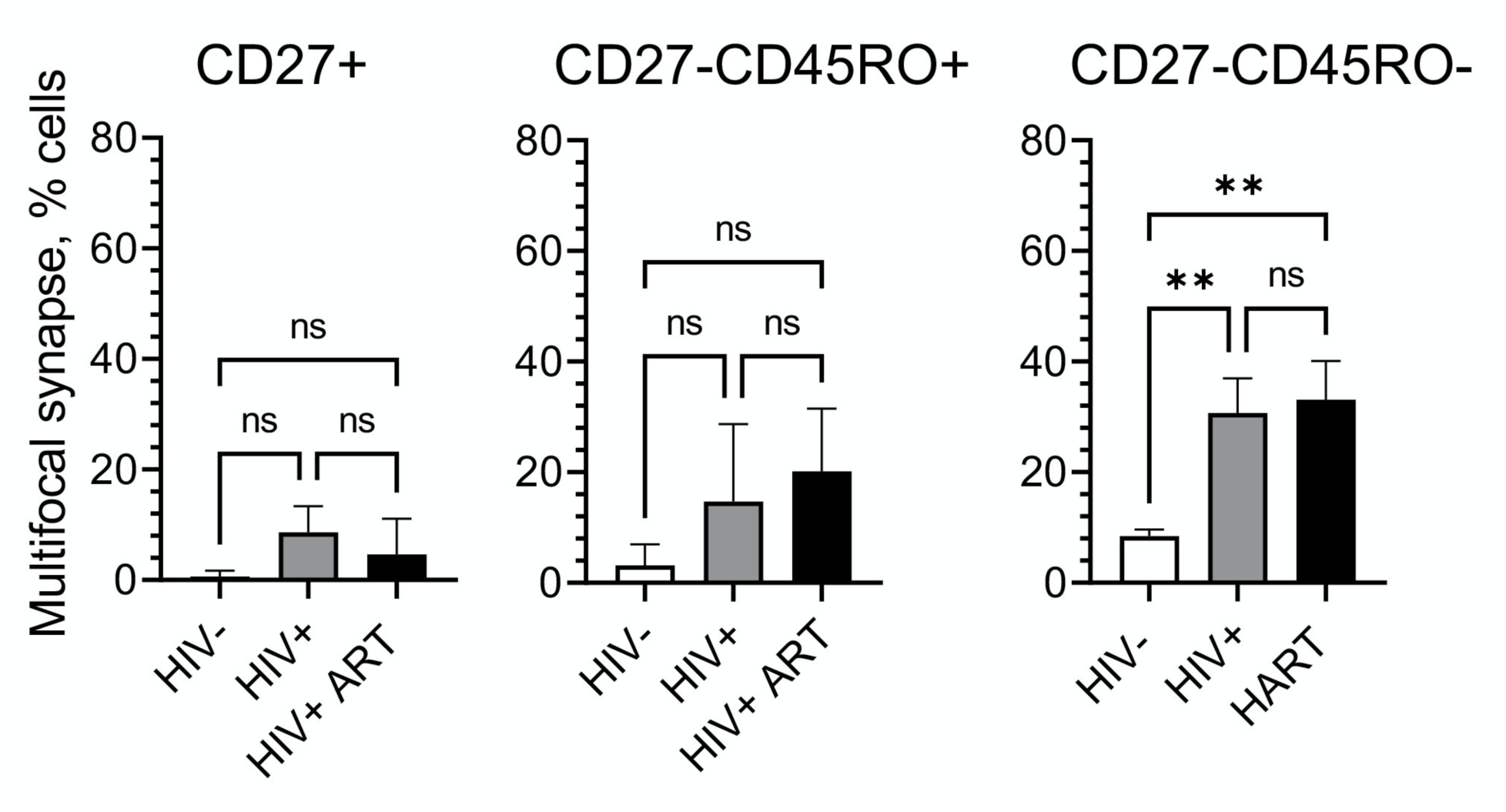
CD8 T cells from HIV+ individuals capable to form noteworthy number of multifocal synapses at late differentiation stage. CD8 T cell subsets from HIV- (**left**), HIV+(**middle**), and ART-treated HIV+ (**right**) individuals were loaded onto bilayer surfaces containing fluorescent labeled anti-CD3 antibodies and ICAM-1 molecules. Interfaces formed between bilayers and T cells were observed with confocal microscopy for 30 min. The data represents mean values (±SD) of three independent experiment for each donor group with different infection status. Statistical significance between subject groups was determined using one-way ANOVA with Tukey’s post-test.

**Supplemental Figure 8.**
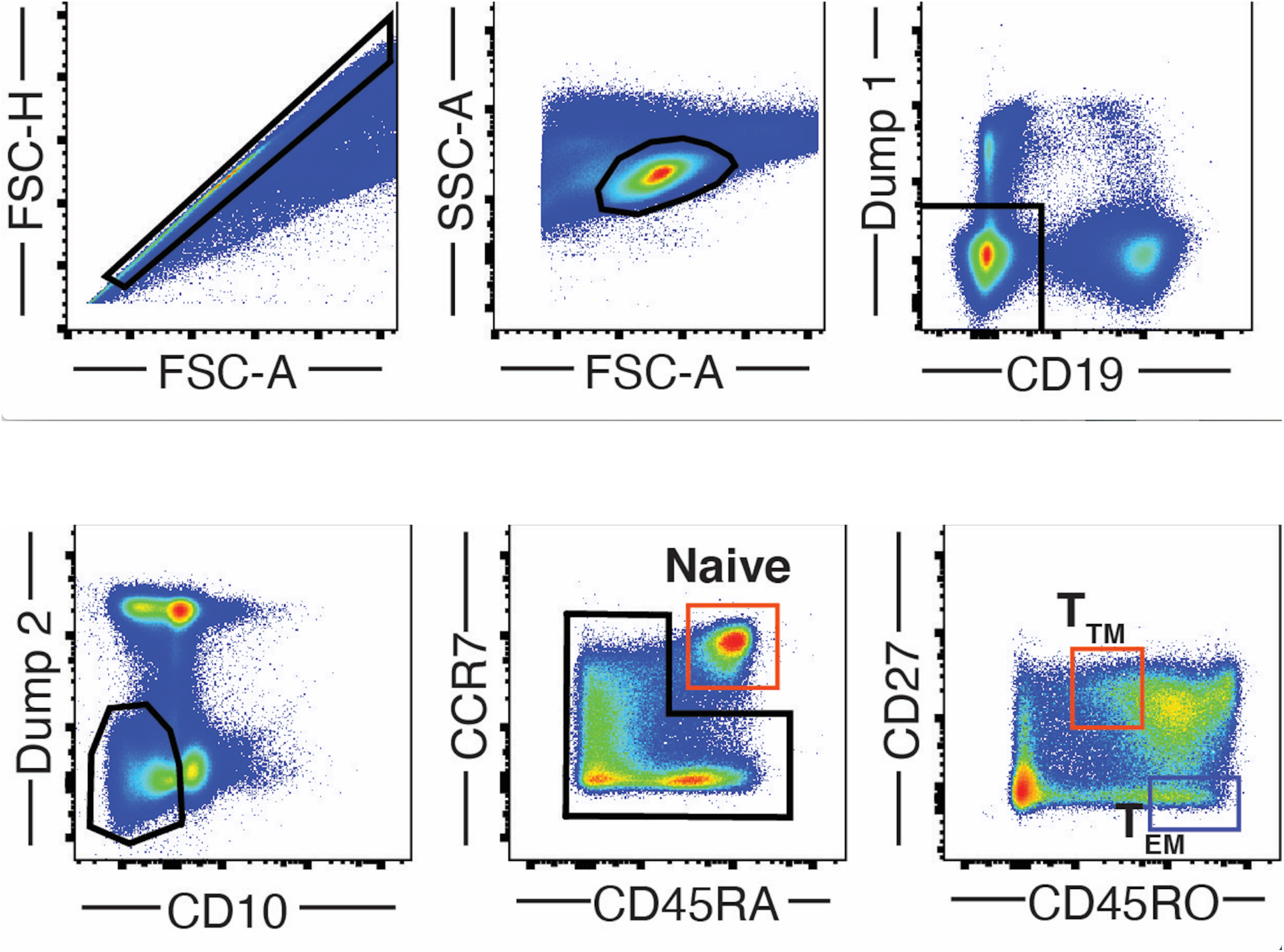
Gating strategy for flow cytometry sorting of CD8+ T cell subsets. Single lymphocytes were first characterized by morphology. Dead cells, CD14+ and CD16+ cells were excluded using Dump 1 (anti-Human CD14, anti-Human CD16, LIVE/DEAD™ Fixable Aqua Dead Cell Stain Kit, see Table 1), and CD19- cells were gated. CD4+ and CD56+ cells were then excluded using Dump 2 (anti-Human CD56, anti-Human CD4, see Table 1), and CD10+ cells were gated out. Naïve cells were characterized as CD45RA+ CCR7+ cells. Transitional (T_TM_, red gate) and effector (T_EM_, blue gate) were identified within the non-naïve cells. T_TM_ were characterized as CD27+ CD45ROlow, and T_EM_ were identified as CD27- CD45RO+ cells. Representative example on an HIV-negative individual.

**Supplemental Figure 9.**
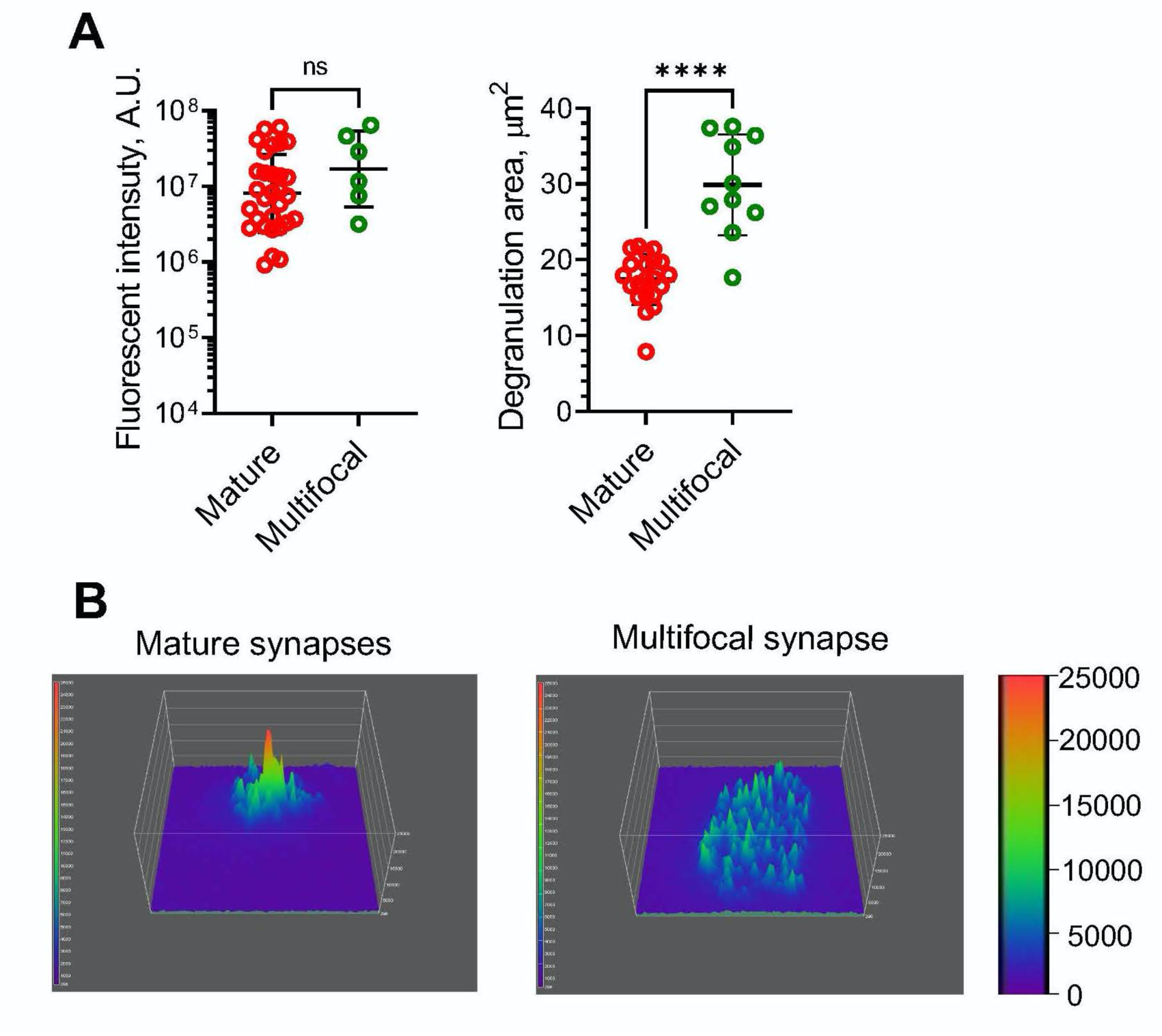
Despite similar amount of released granule, CD8 T cell establishing mature synapses show more efficient delivery (focused vs dispersed) of the cytolytic granules. Effector memory (EM) CD8 T cells from chronic HIV+ donors were isolated by flow cytometry sorting and exposed to bilayer surface presenting fluorescently labeled anti-CD3 antibodies and ICAM-1 in the buffer containing anti-CD107a Fab fragments. The images of interface were taken by TIRF microscope 30 minutes after initial T cell-bilayer contact. (**A**, left panel) CD8 T cells forming mature and multifocal synapses released similar amounts of granules at the interface. CD107a level was measured at mature synapses (red empty circles) and multifocal interfaces (green empty circles). Geometric means and SD are shown as horizontal black lines and black error bars correspondingly; non-parametric two-tailed Mann– Whitney test do not show statistical difference within group. (**A**, right panel) CD8 T cells degranulation were observed on small contact area when the cells form mature but not multifocal synapses. Regions were drawn around areas where degranulation foci were observed, and areas of those regions were determined. Results of one representative experiment is shown (N=3). For each condition, bar represent standard deviation of the mean, n≥10; ****p<0.0001 as established by non-parametric two-tailed Mann–Whitney test. (**B**) Representative surface intensity plot of EM CD8 T cell degranulation at mature synapse (**left**) and multifocal synapse (**right**).

## MOVIE LEGENDS

**Movie 1.** Formation of mature synapse at interface between CD8 T cell and planar bilayer containing fluorescent-labeled anti-CD3 antibodies and ICAM-1. After the cells were exposed to the bilayer, images were taken every 2 minutes for 30 minutes. The movie was assembled using MetaMorph software. (**A**) Quick segregation of ICAM-1 and anti-CD3 molecules that informs about segregation of LFA-1 and TCR/CD3 on the T cell surface, correspondingly, and formation of stable pSMAC and cSMAC zones are noted. Movie represents the entire stack of confocal and IRM images; anti-CD3 antibodies are in green, ICAM-1 molecules are in blue, IRM images are in red. (**B**) After short period of movement and initial contact with the bilayer interface, the cell position over bilayer surface is stabilized. Movie represents stack of bright field images of the same cell as in (**A**).

**Movie 2.** Formation of TCR/CD3 focal interface between CD8 T cell and planar bilayer containing fluorescent-labeled anti-CD3 antibodies and ICAM-1. The cells were exposed to the bilayer, and images were taken every 2 minutes for 30 minutes. The movie was montaged with MetaMorph software. (**A**) Formation and merging anti-CD3 antibody clusters is observed. Movie represents the entire stack of confocal and IRM images; anti-CD3 antibodies are in green, ICAM-1 molecules are in blue, IRM images are in red. **B**) Formation of uropod-like structure linked to the bilayer surface is observed at anti-CD3 antibodies accumulation spots. Movie represents stack of bright field images of the same cell as in (**A**).

**Movie 3.** Formation of kinapse at interface between a CD8 T cell and a planar bilayer containing fluorescent-labeled anti-CD3 antibodies and ICAM-1. The cells were exposed to the bilayer, and images were taken every 2 minutes for 30 minutes. The movie was assembled using MetaMorph software. (**A**) After contact with the bilayer, the cell form transient ‘bull-eye’ structure that converts during movement into asymmetric kinapse with dSMAC zone under leading lamellipodium, middle pSMAC zone and accumulated anti-CD3 antibodies in the trailing edge. Movie represents the entire stack of confocal and IRM images; anti-CD3 antibodies are in green, ICAM-1 molecules are in blue, IRM images are in red. (**B**) The cell movement, formation of leading lamellipodium and trailing uropod are observed. Movie represents stack of bright field images of the same cell as in (**A**).

**Movie 4.** Formation of multifocal synapse at interface between a CD8 T cell and a planar bilayer containing fluorescent-labeled anti-CD3 antibodies and ICAM-1. After the cells were exposed to the bilayer, images were taken every 2 minutes for 30 minutes. The movie was montaged with MetaMorph software. (**A**) Clusters of anti-CD3 molecules dispersed among accumulated ICAM-1. Small clusters fused together to form larger clusters, but complete segregation was not achieved for the period of observation. Movie represents the entire stack of confocal and IRM images; anti-CD3 antibodies are in green, ICAM-1 molecules are in blue, IRM images are in red. (**B**) After contact with the bilayer, the cell forms lamellipodium structure, which increase T cell contact area during immunological synapse formation. Movie represents stack of bright field images of the same cell as in (**A**).

**Movie 5.** Degranulation pattern of CD8 T cells that form mature synapse on bilayer surface containing fluorescent-labeled anti-CD3 antibodies and ICAM-1 in the presence of soluble fluorescent-labeled anti-CD107a Fab fragments. CD8 T cells were allowed to adhere to the bilayer for 4 minutes, and then the images were taken every minute for the entire time of observation. The images were montage into the movie using Metamorph software. (**A**) TIRF microscopy was used for analysis anti-CD3 (green) and anti-CD107a (red) antibodies distribution and wide-field fluorescent microscopy was applied to assess ICAM-1 accumulation; only granule and ICAM-1 position are shown in the movie (**B**). Vast majority of granule were observed inside the ICAM-1 ring, the location where anti-CD3 antibodies accumulated.

**Movie 6.** Degranulation pattern at TCR/CD3 focal synapses that were formed between CD8 T cell and the bilayer presenting fluorescent-labeled anti-CD3 antibodies and ICAM-1 in the presence of soluble fluorescent-labeled anti-CD107a Fab fragments. CD8 T cells were allowed to form contact with the bilayer during 4 minutes, and then the images were taken at a rate one frame/min. (**A**) Anti-CD3 (green) and anti-CD107a (red) antibodies position were assess using TIRF microscopy, while ICAM-1 (blue) distribution was evaluated with wide-field fluorescent microscopy. The movie was prepared using Metamorph software. Pattern of degranulation and position of ICAM-1 and anti-CD3 antibodies are filmed. (**B**) Degranulation was observed around large anti-CD3 antibodies spot and fades with time. Only granule and ICAM-1 position are shown.

**Movie 7.** Degranulation of CD8 T cell that forms kinapse and is moving across bilayer surface. (**A**) TIRF microscopy used for analysis anti-CD3 (green) and anti-CD107a (red) antibodies distribution and wide-field fluorescent microscopy were applied for assess ICAM-1 (blue) accumulation. CD8 T cells were loaded together with anti-CD107a Fab fragments on the bilayer containing fluorescent-labeled anti-CD3 antibodies and ICAM-1 molecules. After 4 minutes, the images were taken every minute for the entire time of observation and then montaged into the movie using Metamorph software. The cell transiently forms classical bull eye structure with granule inside of ICAM-1 ring. Then the synapse loses the symmetry, and cell started to move to form kinapse structure at the interface. At this stage, the granule release observed mostly in or nearby the area of ICAM-1 accumulation. (**B**) Only granule and ICAM-1 distribution are shown.

**Movie 8.** Degranulation pattern of CD8 T cell that forms multifocal synapse. (**A**) TIRF microscopy used to visualize anti-CD3 (green) and anti-CD107a (red) antibodies distribution and wide-field fluorescent microscopy were applied to assess ICAM-1 (blue) accumulation. CD8 T cells were loaded together with anti-CD107a Fab fragments on the bilayer containing fluorescent-labeled anti-CD3 antibodies and ICAM-1 molecules. After 4 minutes, the images were taken every minute for the entire time of observation and then montaged into the movie using Metamorph software. The granule release occurs across entire cell-bilayer interface in multiple locations, which are mostly devoid of ICAM-1. (**B**) Only granule and ICAM-1 distribution are shown.

